# A chromosome-level genome assembly of *Solanum chilense*, a tomato wild relative associated with resistance to salinity and drought

**DOI:** 10.1101/2023.11.17.567531

**Authors:** Corentin Molitor, Tomasz J. Kurowski, Pedro M Fidalgo de Almeida, Zoltan Kevei, Daniel J. Spindlow, Steffimol R. Chacko Kaitholil, Justice U. Iheanyichi, HC Prasanna, Andrew J. Thompson, Fady R. Mohareb

## Abstract

*Solanum chilense* is a wild relative of tomato reported to exhibit resistance to biotic and abiotic stresses. There is potential to improve tomato cultivars via breeding with wild relatives, a process greatly accelerated by suitable genomic and genetic resources. In this study we generated a high-quality, chromosome-level, *de novo* assembly for the *S. chilense* accession LA1972 using a hybrid assembly strategy with ∼180 Gbp of Illumina short reads and ∼50 Gbp long PacBio reads. Further scaffolding was performed using Bionano optical maps and 10x Chromium® reads. The resulting sequences were arranged into 12 pseudomolecules using Hi-C sequencing. This resulted in a 901 Mbp assembly, with a completeness of 95%, as determined by Benchmarking with Universal Single-Copy Orthologs (BUSCO). Sequencing of RNA from multiple tissues resulting in ∼219 Gbp of reads was used to annotate the genome assembly with an RNA-Seq guided gene prediction, and for a *de novo* transcriptome assembly. This chromosome-level, high-quality reference genome for *S. chilense* accession LA1972 will support future breeding efforts for more sustainable tomato production. Gene sequences related to drought and salt resistance were compared between *S. chilense* and *S. lycopersicum* to identify amino acid variations with high potential for functional impact. These variants were subsequently analysed in 84 resequenced tomato lines across 12 different related species to explore the variant distributions. We identified a set of 7 putative impactful amino acid variants some of which may also impact on fruit development for example the e*thylene-responsive transcription factor WIN1* and *ethylene-insensitive protein 2*. These variants could be tested for their ability to confer functional phenotypes to cultivars that have lost these variants.

## Introduction

Non-starchy vegetables are one of the cornerstones of a healthy human diet [1]. Domesticated tomato (*Solanum lycopersicum* <u>L.</u>) is the most-consumed non-starchy vegetable in the world, reaching 180 million tonnes of production in 2019, equivalent to every human eating one 63g tomato every day of the year (FAO, 2021). Tomato breeding to maintain and enhance yield, resilience, sustainability, and nutrition of tomato crops is therefore an important endeavour. There has been concern that modern breeding is leading to a reduction in genetic diversity, leading to less resilience against shifting pest and disease risks and lower human nutrition in favour of yield. At least in The Netherlands this was a temporary problem that started to reverse in the 1970s, with an eight fold increase in genetic diversity that delivered greater disease resistance and improved fruit quality via the introduction of DNA from wild species into cultivated tomato (Schouten et al., 2019).

The tomato reference genome [2] and short-read resequencing [3] provides a wealth of SNP and InDel polymorphism data across most wild species, whereas long-read platforms have been used to improve assemblies [4] and report variants [5] and structural variants [6] in tomato and its most closely related sub-species and wild species, *S. lycopersicum* var. *cerasiforme, S. pimpinellifolium*, *S. cheesmaniae* and *S. galapagense*. Long-reads have also been used to create a “graph pan genome” [7], but only for cultivated tomato and its most closely related wild species (*S. lycopersicum* var. *cerasiforme* and *S. pimpinellifolium*). A chromosome level assembly is also reported for close relative *S. pimpinellifolium* [8]. For more distantly related species there are a few high-quality, chromosome-level assemblies available: *S. pennellii* (Bolger et al., 2014), *S. sitiens* [9] and *S. lycopersicoides* (Powell et al., 2022). These provide the genomic resources to facilitate gene functional studies and marker discoveries needed for wide introgression breeding.

*S. chilense* is a wild relative of tomato, classified into the *Solanum* section *Lycopersicon* in the Eriopersicon group along with *S. habrochaites*, *S. huaylasense*, *S. corneliomulleri* and *S. peruvianum* (Peralta et al., 2008); population genetic studies estimate that it diverged from *S. peruvianum* less than 0.55 million years ago [10]. It is diploid (n = 12) allogamous and self-incompatible; successful crossing with cultivated tomato is very rare (Rick, 1979), requiring bridging lines or embryo rescue.

It is native to southern Peru and northern Chile where it can grow in altitudes ranging from sea level to over 3,500 meters [11, 12] and is often found “in the extremely dry high-elevation deserts of the western Andean slope…and in the unique lomas habitat” where lomas are “small areas of vegetation occurring as islands in a sea of hyper-arid desert” (Peralta et al., 2008). The species characteristically has greyish pubescent leaves, straight anther tubes with exerted stigma, long erect peduncles and inedible green fruits of ∼1 cm coated in short trichomes when immature and developing a purple stripe when mature (Figure 1). *S. chilense* is considered, based on its ecological distribution, to be resistant to extreme environments, including drought, high salinity and low temperature stresses [11, 13]. One physiological study claims salinity resistance for *S. chilense* LA4107 and its F_1_ hybrid with cultivated tomato [14]. Moreover, *S. chilense* is resistant to pathogens, notably the *Tomato Yellow Leaf Curl Virus,* the *Cucumber Mosaic Virus* [15] and Tomato Mottle Virus [16].

**Figure 1:**
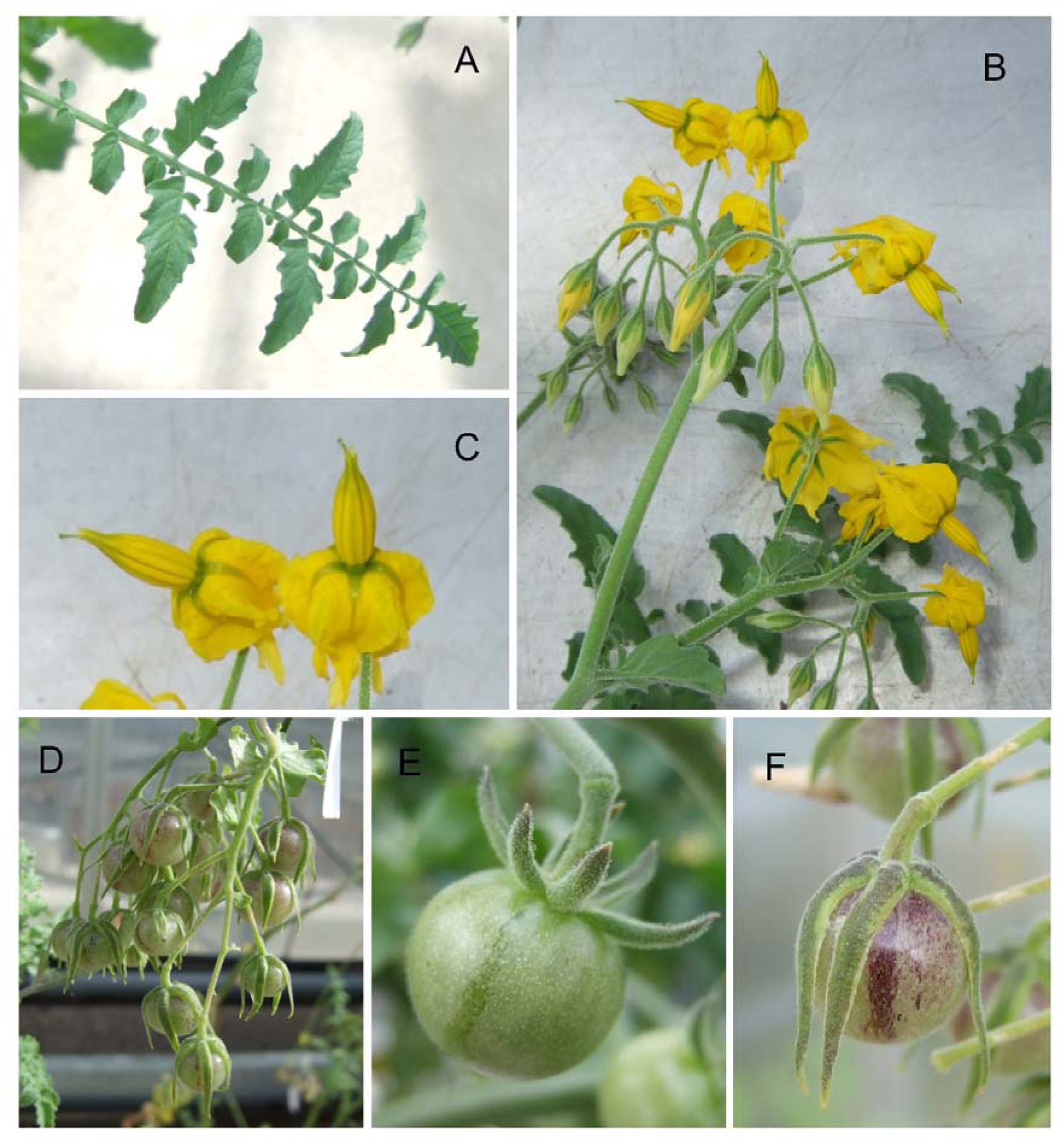
Images of plants of Solanum chilense LA1972 growing in a glasshouse. A: fully expanded leaf; B: inflorescence; C: detail of flowers from image B; D: truss of ripening fruit; E: single fruit at green ripe stage; F: single ripe fruit. Twenty plants were mass-sibling pollinated to achieve fruit set.

Böndel et al. (2015) demonstrated that in the southern range, *S. chilense* shows high genetic variation between the coastal and high-altitude populations, suggesting different local adaptation between the eastern and western sides of the Atacama desert [17]. The high-level of heterozygosity arising from the allogamous, self-incompatible mating system creates a challenge to generate a high-quality reference genome. A scaffold-level assembly for *S. chilense* (accession LA3111) is the only genome assembly reported so far for this species; the data allowed the identification of two unique Coiled-Coil domain containing NLR subfamilies: CNL20 and CNL21 [18].

The aim of this work was to produce a high-quality reference genome for *S. chilense* LA1972 using a hybrid-sequencing strategy including both Illumina short reads and Pacific Biosciences long reads, followed by Bionano optical mapping and Hi-C sequencing to orient, and order the scaffolds into pseudomolecules. Additionally, the generated genome assembly has been complemented with a high-quality and functionally annotated *de novo* transcriptome assembly with gene models. Our assembly provides a resource that will underpin the introgression of beneficial traits into cultivated tomato.

We chose *S. chilense* LA1972 because it was collected from an extremely dry environment and is classified as “drought tolerant” in the C.M. Rick, Tomato Genetics Resource Centre (TGRC) catalogue, and because of the availability of successful embryo-rescued progeny from crossing with cultivated tomato for development of genetic resources. Although leaves or whole plants of *S. pennellii* are more resistant to desiccation compared to cultivated tomato, the same is not considered to be true for *S. chilense*, where it was hypothesised that its drought resistance is due to the foraging ability of its root system in rocky riverbeds, rather than leaf traits [19].

## Materials and Methods

### Plant materials

*S. chilense* LA1972 seeds were obtained from the UC Davis/C.M. Rick Tomato Genetics Resource Centre (TGRC) maintained by the Department of Plant Sciences, University of California, Davis, CA 95616, USA. LA3111 from which the scaffold level assembly was reported [20] was collected from Tarata, Tacna province at elevation 3070 m; LA1972 was collected from Rio Sama, also Tacna province at elevation 650 m; these two locations are approximately 70 km apart.

Twenty plants were raised and crossed between siblings to bulk seeds for physiological analysis, but one individual plant was used for all DNA and RNA extractions – this plant is maintained through clonal propagation at Cranfield University, UK. Seed from *S. lycopersicum* cv. Kashi Amrit was obtained from the Division of Crop Improvement, ICAR-Indian Institute of Vegetable Research, Varanasi, India.

### DNA and RNA extraction

*S. chilense* leaves used for DNA and RNA extraction were obtained from plant grown to flowering stage in a glasshouse facility at Cranfield University, UK. The DNeasy and RNeasy Plant Mini Kits (Qiagen, Manchester, UK) were used to prepare genomic DNA and total RNA for Illumina sequencing according to the manufacturer’s instructions. High molecular weight (HMW) genomic DNA was extracted for PacBio sequencing and Bionano optical mapping. The HMW DNA was prepared by the Earlham Institute, Norwich, UK using the Bionano Prep^TM^ Plant Tissue DNA isolation kit.

### Sequencing data

Two PCR-free, paired-end Illumina libraries were prepared and sequenced on an Illumina HiSeq2500^TM^ platform at the Earlham Institute (UK), with a read length of 250 base pairs (bp) and a mean insert size of 395 bp. The whole genome sequencing yielded a total of ∼180 Gbp and the quality of the reads was assessed with FASTQC v0.11.

Pacific Bioscience long reads were obtained from two different platforms, namely RS-II and Sequel. Using RS-II, ∼16 Gbp of data were generated in 2,186,914 reads, with a N50 of 9,384 bp. The longest read was 49,532 bp long. For the Sequel platform, ∼34 Gbp of data were generated in 3,818,160 reads, with a N50 of 14,770 bp. The longest read was 160,787 bp long. For each platform, the bam files were converted into a multi-sequence fasta file, containing all the reads. The two resulting fasta files were concatenated into one, which was used in the subsequent assembly steps.

Optical maps were generated with the BioNano Irys platform, at the Earlham Institute, yielding ∼314 Gbp of molecules larger than 100 kbp. The *Bss*SI restriction enzyme was used and resulted in a label density of ∼11 per 100 kbp.

A single library of Paired-End 10X Chromium, generating 28 Gbp of data, was sequenced at the Earlham Institute following 10x Genomics guidelines for genomes between 0.1 and 1.6 Gbp. The fastq files were processed with the “basic” pipeline from LongRanger v2.2.2 which interleaved the two fastq files and performed quality control (read trimming, barcode error correction and barcode whitelisting). The 10x molecule barcode, present in the first 16 bp of each left read, was removed and added to the corresponding read pair identifiers, which is required for most downstream analyses. Finally, the Arcs pipeline [21], used as part of the assembly process, requires the 10x reads to have the barcode as part of their name: this was achieved with a custom Perl script “10x_custom_script.pl” (see Data availability). Reads without a barcode were removed, which corresponded to ∼17 million reads, representing 5% of the total number.

Finally, chromosome conformation Hi-C data was generated using the Arima-HiC kit, according to the manufacturer’s protocol (Arima Genomics, San Diego, US). A Hi-C library was prepared using the Arima approach, which uses a mixture of restriction enzymes cutting chromatin at the following sequence motifs: ^GATC, G^ANTC, G^TNA and T^TAA. The library was sequenced on an Illumina HiSeq X^TM^ platform, which yielded a total of 367,560,717 read pairs (2 x 150 bp). The quality of the library preparation was assessed by Arima Genomics using human control GM cells which identified ∼56% of long-range *cis* interactions, ∼24% of *trans* interactions and 0.2% duplication.

For gene models prediction and functional annotation, 15 Illumina 125 bp paired-end RNA-Seq libraries were sequenced (HiSeq2500^TM^) generating 219 Gbp of data. The 15 tissue samples were: 2 x fruit and sepals at 14 days after pollination (dap); fruit at 36 dap; sepals at 36 dap; fruit at 46; root with or without dehydration treatment; fully expanded leaf with or without dehydration treatment; 2 x flower; 2 x stem; senescing leaf; young expanding leaf and meristem combined. Deyhdration treatments were achieved by drying tissue on the laboratory bench under ambient conditions until 10% of fresh weight was lost (inducing loss of turgor except in stem).

The quality of the RNA-Seq reads was assessed with FastQC. The correction of erroneous K-mers was performed using RCorrector v1.0.3.1 [22], a tool which utilises a K-mer spectrum based method to convert rare K-mers (a K-mer size of 19 was used) into those that are more commonly found within the assembly. Reads deemed unfixable by RCorrector were removed using FilterUncorrectablePEfastq.py from the TranscriptomeAssemblyTools package. Bases with a PHRED score < 5 were trimmed and adapter sequences and trimmed reads below a length of 100 bases were removed using Trim Galore [23], a wrapper around Cutadapt [24].

### Genome size estimation

The size of the genome was estimated by performing a a k-mer based analysis using the Illumina short reads [25]: 25-mers from the two Illumina libraries were counted with Jellyfish v2.2.3 [26] with the -C parameter to consider both strands. A total of 133 billion 25-mers were counted and plotted as a histogram (Figure 3). The 18 billion 25-mers with an occurrence lower than 30 were considered artifacts (as they probably spawned from sequencing errors) and were disregarded during the genome size estimation.

### *De novo* assembly strategy

The hybrid assembler MaSuRCA v3.2.7 [27], generated the *de novo* contig assembly, based on both the paired-end Illumina reads and the combined RS-II and Sequel PacBio reads. The k-mer size of 127 was automatically determined by MaSuRCA, the ploidy was set to 2 and the k-mer count threshold was set to 2 as the Illumina coverage was more than 100x. The Jellyfish hash size was set to 125 billion and 64 threads were used to speed up computation time; the default values were kept for the remaining parameters.

Despite the satisfactory contiguity, the number of artifact duplications was high, as assessed by BUSCO [28] and resulted in larger than expected total genome size (See the Genome size estimation section). This is expected when attempting to assemble a heterozygous, out-breeding, wild species as it is the case here. Redundans v0.13a [29], was applied to the contig assembly with the Illumina reads, in order to remove artifact duplications and then scaffold the resulting reduced assembly. Next, SSPACE v1-1 [30], further scaffolded the assembly, using the default parameters and the combined PacBio long reads (RSII and Sequel).

Optical maps from BioNano Genomics, obtained with the BssSI (GACGAG) restriction enzyme served as input to the “Hybrid Scaffold” pipeline, which super-scaffolded the assembly. The conflict filter levels (options *-B* and *-N*) were set to 1 and the xml file describing the remaining parameters (option *-c*) is available at https://github.com/MCorentin/Solanum_chilense_assembly. The restriction enzyme used in this analysis was manually added to Hybrid Scaffold, as it is not supported by default. This conservative tool removes all unmapped scaffolds from the assembly, which reduces the completeness, hence the Perl script “hybridScaffold_finish_fasta.pl” [31] was run to reintegrate the discarded scaffolds into the assembly.

The super-scaffolded assembly was given as input to Arcs v8.25 [21] and LINKS v1.8.6 pipeline [32], which uses long range information from the 10x Chromium reads in order to further scaffold the assembly. First, the interleaved 10x reads were aligned to the super-scaffolded assembly with bwa v0.7.17 [33] using the “mem” algorithm. Then a Graphviz Dot file, representing scaffolds as nodes and evidence that two scaffolds are linked as edges, was generated with Arcs. The following parameters were chosen: the minimum sequence identity for read alignment was set to 95% (option -s), the range for the barcode multiplicity was set to 30-10000 (option-m) and default values were kept for the remaining parameters. The *makeTSVfile.py* python script translated the Graphviz Dot file to a tsv file, which contains all possible oriented sequence pairs with the number of supporting barcodes. This tsv file was given as input to LINKS, with default parameters except for the k-mer size, which was set to 20, to generate the super-scaffolded fasta file.

Assembly polishing was performed via two iterations of Pilon v1.22 [34]. For each iteration, first the Illumina short reads were aligned to the super-scaffolded assembly with *bwa mem*, then the resulting SAM file was converted to a BAM file, sorted and indexed with SAMtools v1.9 [35]. Pilon was run on the assembly fasta file with the aligned BAM files and the following parameters: *--changes*, to generate a log file listing all the changes and *--fix all*, to fix individual base errors, small InDels (insertion/deletion), gap sizes and local misassemblies.

The polished assembly was inputted to BBmap’s *dedupe.sh* script v37.72 to remove duplicated sequences from the assembly, based on sequence similarity. For this step, *storequality* was set to false; *absorbrc* was set to true to absorb reverse complements as well as normal orientation; *touppercase* was set to true to avoid mismatches due to lowercases; *minidentity*, representing the minimum sequence similarity to consider two sequences as duplicated was set to 90%; *minlengthpercent* and *minoverlappercent* were both set to 0 to ignore filtering based on contig lengths and overlap; the maximum number of allowed substitutions, *maxsubs*, and InDels, *maxedits* were set to 40,000 and 1,000 respectively; and finally the seed length, *k*, was set to 31. The values for *maxsubs* and *maxedits* were chosen empirically after testing a range of different values and assessing the resulting assembly with Quast and BUSCO (See Supplementary Table 9).

The final step of the scaffolding was done with GapFiller v1-10 [36], which harnessed information from the paired-end Illumina reads to resize and fill gaps between or within scaffolds. The minimum number of overlapping bases with the edge of the gap was set to 30 (option -m), and default values were kept for the remaining parameters, notably the number of iterations which was set to 10.

An overview of the whole assembly pipeline is available as Figure 2.

**Figure 2:**
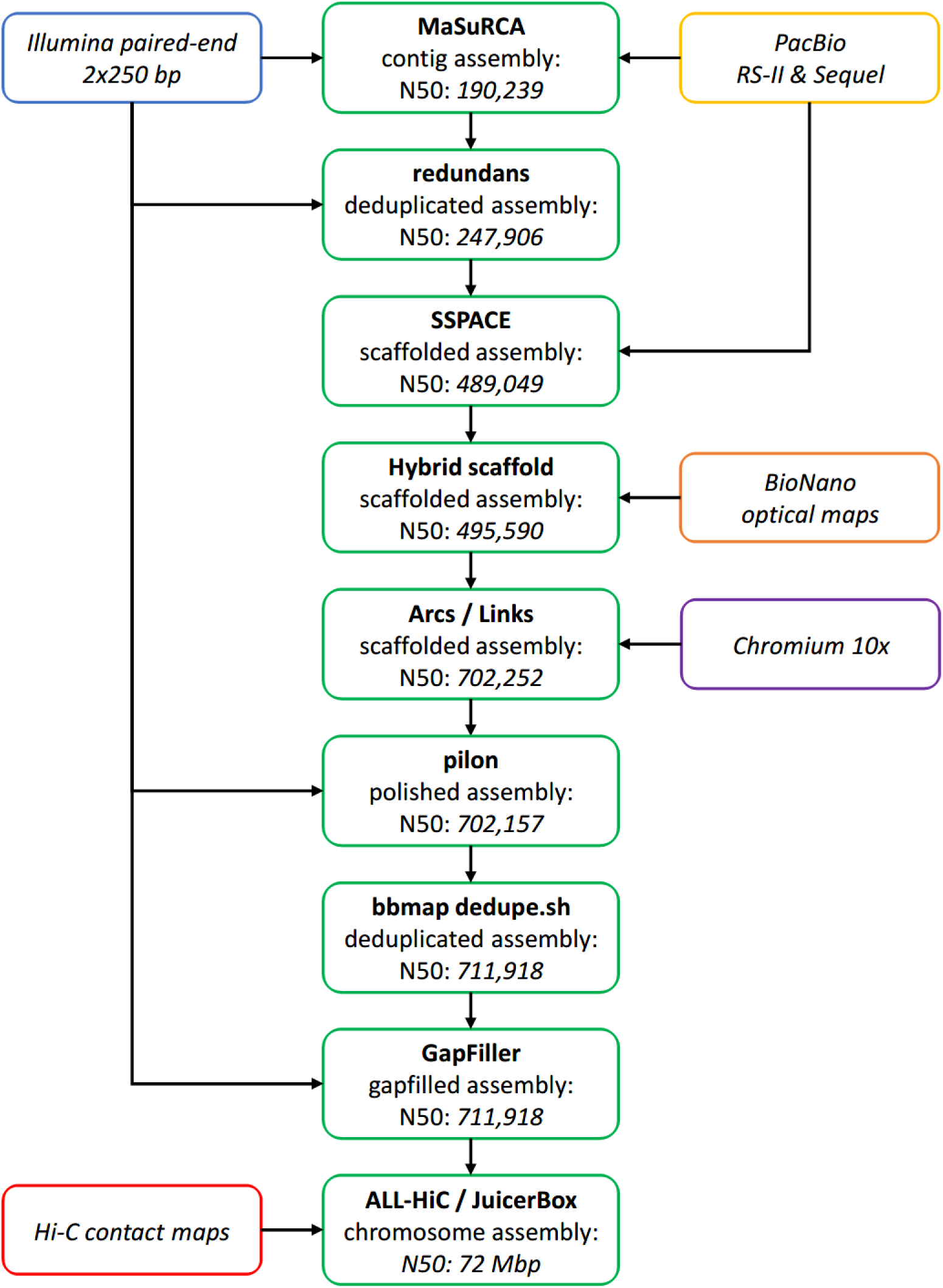
Overview of the assembly pipeline. Assembly steps are represented in green, with the resulting N50 specified inside the box (in basepairs, unless stated otherwise).

### Construction of pseudomolecules from the Hi-C data

Contact information, obtained from mapping the Hi-C reads against the assembly, was used to orient and order the scaffolds into chromosome-sized sequences. First, the Hi-C reads were trimmed with Trimmomatic v0.39 [37] using a sliding window of 4 bases, and trimming the reads when the average base quality reached below 20 (SLIDINGWINDOW:4:20), as well as removing reads smaller than 50 bases (MINLEN:50). The trimmed reads were mapped to the gapfilled assembly with bwa v0.7.17 using the “*-SP*” options to align the pairs as independent single-end reads, while keeping all the appropriate pair-related flags in the resulting SAM file, and the “*-5*” option to only keep alignments with the smallest coordinates as primary, when dealing with split alignments. This last option is beneficial when aligning Hi-C data, as it reduces the number of secondary mappings and the “noise” from the Hi-C alignment. The resulting SAM file was filtered with SAMtools v1.9 [35] to remove reads in non-primary, supplementary and unpaired alignments. The SAM file was further processed with the *PreprocessSAMs.pl* script from LACHESIS [38] to remove redundant, chimeric and uninformative read pairs, while retaining significant Hi-C links from the alignment. This step reduces the file size, and subsequently the I/O time needed to process it.

The ALL-HiC pipeline v0.9.13 [39] oriented and ordered the scaffolds in the assembly. First, the Hi-C reads were aligned against the corrected assembly, using the same parameters (*-SP* and *-5*) and post-processing the SAM file with the same options as before, but with an additional step of removing the alignments with a quality lower than 40 (*samtools view -q 40*). Then, the *ALLHiC_partition* script clustered the scaffolds into 12 groups (*-k 12*) as the number of expected chromosomes in the *S. chilense* genome. After the partition step, the *extract* step created ChromLinkMatrix files containing the intrachromosomal links data for each cluster. These ChromLinkMatrix files were given as input to the *optimize* step to find the orientation and ordering best supported by the Hi-C alignments.

We identified some misjoins in our pseudomolecules, based on comparisons against chromosomes from closely related species, namely *Solanum lycopersicum* [40] and *Solanum pennellii* [41]. Juicebox v1.11.08 [42] was used to manually curate the orientation and order of the scaffolds in these regions. First, the *agp* file obtained from ALL-HiC was converted to an *assembly* format with the *agp2assembly.py* script from *phasegenomics* (*Juicebox Utilities*, 2018/2021), then a *hic* file was created using *matlock*, from the alignment of the Hi-C reads against the corrected genome. Finally, the reviewed chromosome-level assembly was converted back into a fasta file with the *juicebox_assembly_converter.py* script.

### Quality assessment

An important step in the generation of a *de novo* reference genome is the quality assessment of the final assembly. Assembly quality metrics were calculated with Quast v4.5 [43]. Completeness and duplication levels were measured with BUSCO v5.3.2 [28] against the OrthoDB *Solanaceae* v10 [44], containing 3,052 highly conserved orthologues from this family. Sequence similarity with closely related species, *S. lycopersicum v4.0* and *S. pennellii* was assessed with Mummer v4.0.0 [45] using the *–mum* and *-c 1000* parameters, to remove noise from the global alignment. Finally, the K-mer Analysis Toolkit (KAT) v2.4.0 [46] assessed the completeness of the assembly by comparing 27-mers obtained from the assembly against those obtained from the Illumina reads.

### Gene prediction and annotation

Genes were predicted from the final assembly using Augustus v3.3 [47] with hints obtained via the alignment of the RNA-Seq reads against the assembly. First, repeats present in the final assembly were masked with RepeatMasker version open-4.0.9 [48] using the *repeats_master.fasta* library of repeats for *Solanum lycopersicum* (obtained from SolGenomics [49]). The *--xsmall* parameter was used to return repetitive regions in lowercases, rather than mask them, which would hinder gene prediction.

Then, the RNA-Seq reads were aligned to the masked assembly with STAR v2.6.0c [50]. Both libraries were aligned with default parameters. The two resulting BAM files were merged with SAMtools and then sorted by query with *samtools sort -n*. The *filterBam* script from Augustus was applied to the sorted BAM file with the *--uniq* and *--paired* parameters to remove the background noise from the alignment. Finally, the hints file, containing information about introns, was generated with the *bam2hints* script from Augustus, using the aforementioned BAM file as input.

Gene prediction was performed with Augustus, which was run with the following parameters: *tomato* was chosen as the species, *softMasking* was set to *on*, to indicate that the assembly was soft-masked, *allow_hinted_splicesites* was set to *atac*, to allow Augustus to predict the rare introns that start with AT and end with AC, and *–alternatives-from-evidence* was set to true to allow the prediction of alternative splicing. The default configuration file for the extrinsic evidence, which lists the used sources for the hints and their “boni” and “mali”, was replaced with *–extrinsicCfgFile*.

The genes predicted with Augustus were annotated using OmicsBox v1.3.11 [51]. The amino acid sequences were blasted against the NCBI-nr database using the *blastp* algorithm, using the following parameters: the *expectation p-value* was set to 1.0e^-3^, the *word size* was set to 5, the *HSP length cutoff* was set to 15 and the *low complexity filter* was turned on. Gene Ontology (GO) mapping and annotation were also performed by OmicsBox based on the blast results. An InterProScan [52] search against all available databases was performed on the FASTA sequences with OmicsBox via the web service offered by the EBI.

### Organelles assemblies and annotations

The assembly of *S. chilense* chloroplast and mitochondrial genomes is described in the supplementary materials.

### *De novo* transcriptome assembly

A *de novo* transcriptome assembly was generated from the RNA-Seq reads using Trinity v2.8.5 [53] with a k-mer size of 25. *In silico* normalization was performed by setting the maximum reads coverage to 50x to speed up the process. After completion of the transcriptome assembly, the redundancy was reduced by clustering similar transcripts with CD-HIT-EST v4.8.1 [54] using a word size of 10 and a sequence identity threshold of 0.95. To remove sequencing artefacts presenting as lowly expressed transcripts, abundance estimation was performed using Trinity’s *align_and_estimate_abundance.pl* and *abundance_estimates_to_matrix.pl* scripts; ultimately a threshold of 1 TPM was selected, and corresponding transcripts were filtered with Trinity’s *filter_low_expr_transcripts.pl* script.

As for the main assembly, completeness of the transcriptome was assessed at each stage throughout the comparison of orthologues within the assembly to the *Solanaceae* orthoDB dataset using BUSCO. Additionally, completeness was further assessed by realigning the RNA-Seq reads back to the assembly using Bowtie2 v2.3.3.1 [55]. Indeed, assembled transcripts may not fully represent the RNA-Seq reads from which they are derived from and thus alignments from properly and improperly paired reads were captured to quantify read representation.

### Transcriptome functional annotation

The final transcriptome assembly was blasted using *blastx-fast* [56] with default parameters against NCBI’s non-redundant proteins database (NR) and manually generated databases from *S. lycopersicum* (*ITAG3.2*), *S. pennellii* (*Spenn-v2-aa- annot.fa*) annotated proteins, obtained via the SolGenomics website, and The Arabidopsis Information Resources’ annotated protein list (*TAIR10*) [57].

The resultant top 20 Blast hits were loaded into OmicsBox v1.3.11, with a HSP cut- off of 33. GO mapping was performed after which annotation was run with a cut-off length if 55, a GO weighting of 5 and e-value filter of 1×10^-6^. Enzyme code mapping was performed for the identification of enzyme codes based upon the GO IDs. Finally, an InterProScan search was done on the assembly to detect GO terms based upon protein signatures.

### Comparative genomics

A comparative genomics analysis was performed on 84 accessions across 12 tomato species focusing on genes related to drought and salt response. The rationale behind this analysis was to identify variants between *S. lycopersicum* and *S. chilense,* and further wild relatives, which could help us to understand potential differences in drought and salt stress resistance, with the hypothesis that these some of these traits had been lost during domestication of *S. lycopersicum*.

First, genes related to drought and salt response were selected, using a set of 16 Gene Ontology (GO) terms, listed in Table 1. The terms were obtained from searching the following keywords, *salt, salinity, water* and *drought*, in the annotated gene list of our *S. chilense* assembly (see: Gene prediction and annotation).

**Table 1:**
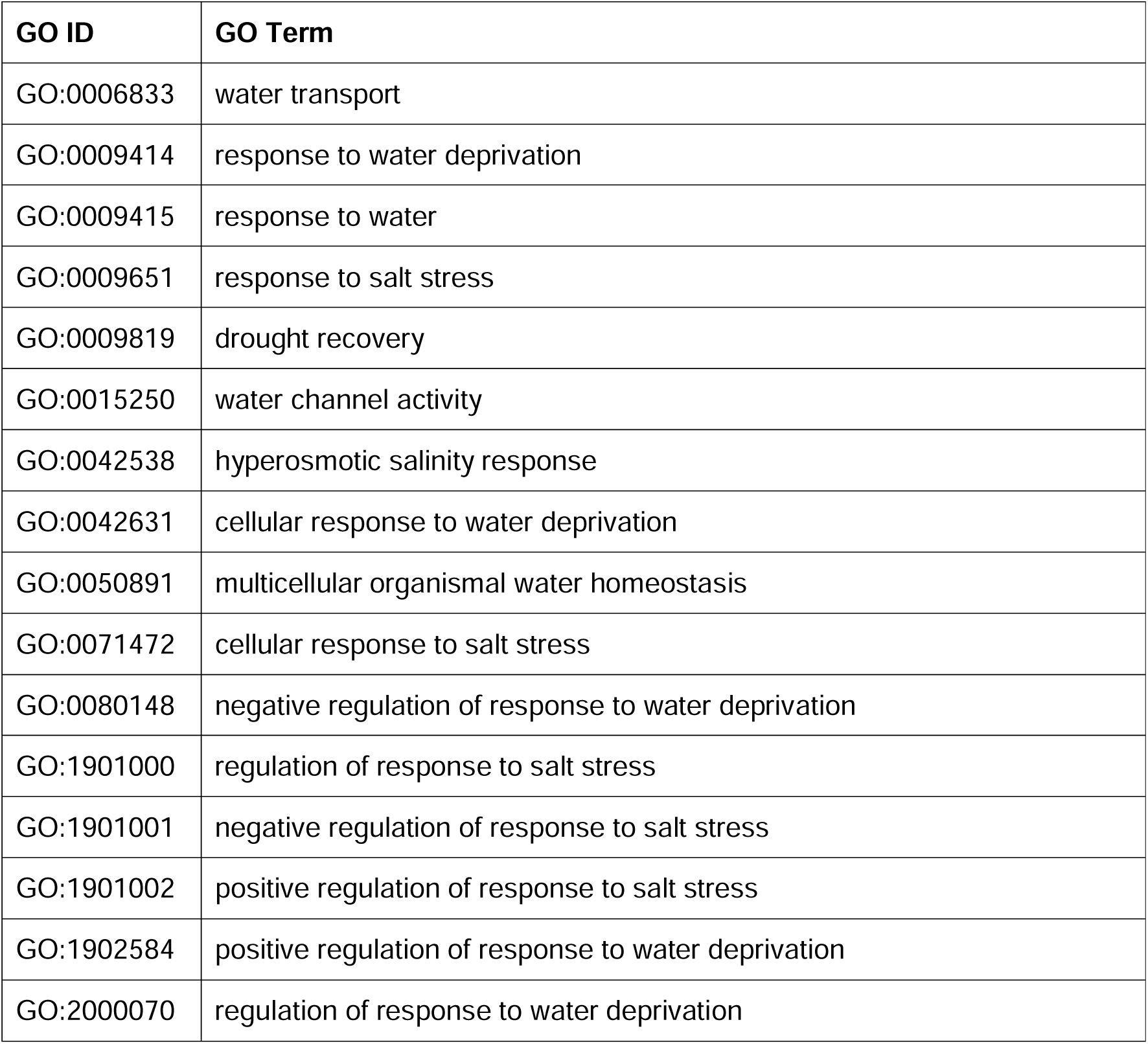
List of Gene Ontology terms related to water and salt response, from the S. chilense annotation, obtained by keyword search.

| GO ID | GO Term |
| --- | --- |
| GO:0006833 | water transport |
| GO:0009414 | response to water deprivation |
| GO:0009415 | response to water |
| GO:0009651 | response to salt stress |
| GO:0009819 | drought recovery |
| GO:0015250 | water channel activity |
| GO:0042538 | hyperosmotic salinity response |
| GO:0042631 | cellular response to water deprivation |
| GO:0050891 | multicellular organismal water homeostasis |
| GO:0071472 | cellular response to salt stress |
| GO:0080148 | negative regulation of response to water deprivation |
| GO:1901000 | regulation of response to salt stress |
| GO:1901001 | negative regulation of response to salt stress |
| GO:1901002 | positive regulation of response to salt stress |
| GO:1902584 | positive regulation of response to water deprivation |
| GO:2000070 | regulation of response to water deprivation |

The sequence of the annotated genes with the aforementioned GO terms in *S. chilense* were extracted with SeqKit v2.3.0 [58]. Orthologues in *S. lycopersicum* were identified via a blast search using blast+ v2.13.0, with the blastn algorithm and the *qcov_hsp_perc* parameter set to 90. Multiple sequence alignments of the protein sequences were performed with MAFFT v7.490 [59]. An in-house Python script was used to extract the amino acid substitutions, insertions, and deletions from the multiple sequence alignment files.

Finally, PROVEAN (PROtein Variant Effect ANalyzer) v1.1.5 [60], SIFT4G [61] and PPVED [62] predicted whether an amino acid substitution affected protein function. The PROVEAN analysis was based on blast+ v2.4.0, the nr database v2.4 and CD-HIT v4.6.1 [54]. The SIFT4G analysis was used with UNIPROT’s uniref90 as the database.

## Results & Discussion

### Genome size estimation

Figure 3 represents the k-mer spectra, plotted as a histogram. The number of remaining 25-mers were divided by the expected homozygous coverage, of 136, as determined by the location of the homozygous peak, and revealed an estimated genome size of 845 Mbp. However, the high heterozygous sequence of the sample might have impacted the accuracy of this result. Detailed statistics about the genome size estimation can be found in Supplementary Table 7.

**Figure 3:**
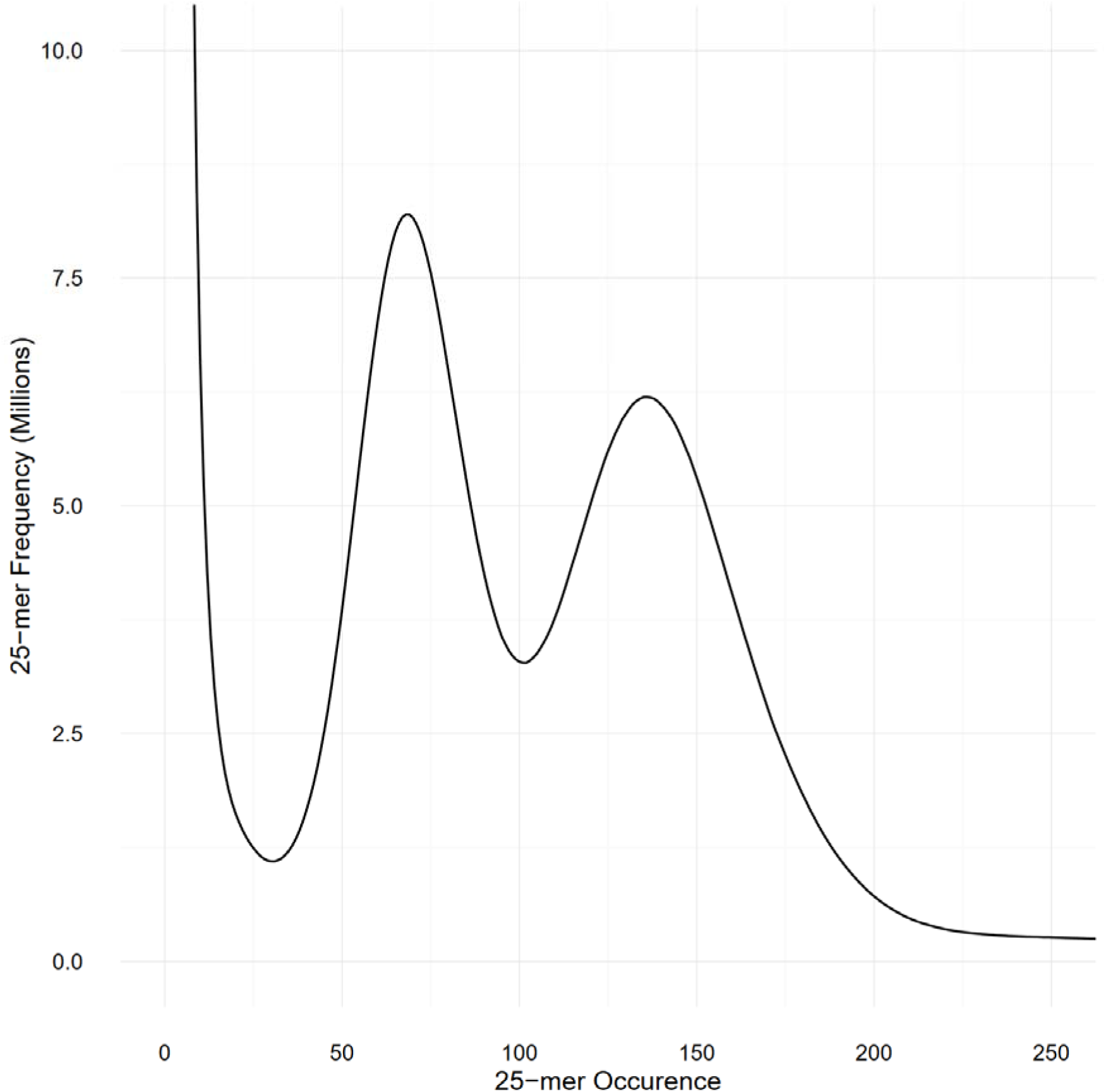
Histogram representing the frequency of the 25-mers present in both Illumina libraries, as counted by Jellyfish. The two peaks, at occurrence 68 and 136 represents respectively, the heterozygous and homozygous peaks and demonstrate the high heterozygosity present in our S. chilense sample.

### The genome assembly

The assembly statistics, as measured by Quast, were computed at each step of the pipeline and the results are available in Table 2. Detailed statistics are available in Supplementary Table 1. The final assembly was also represented as a Circos plot (Figure 4).

**Figure 4:**
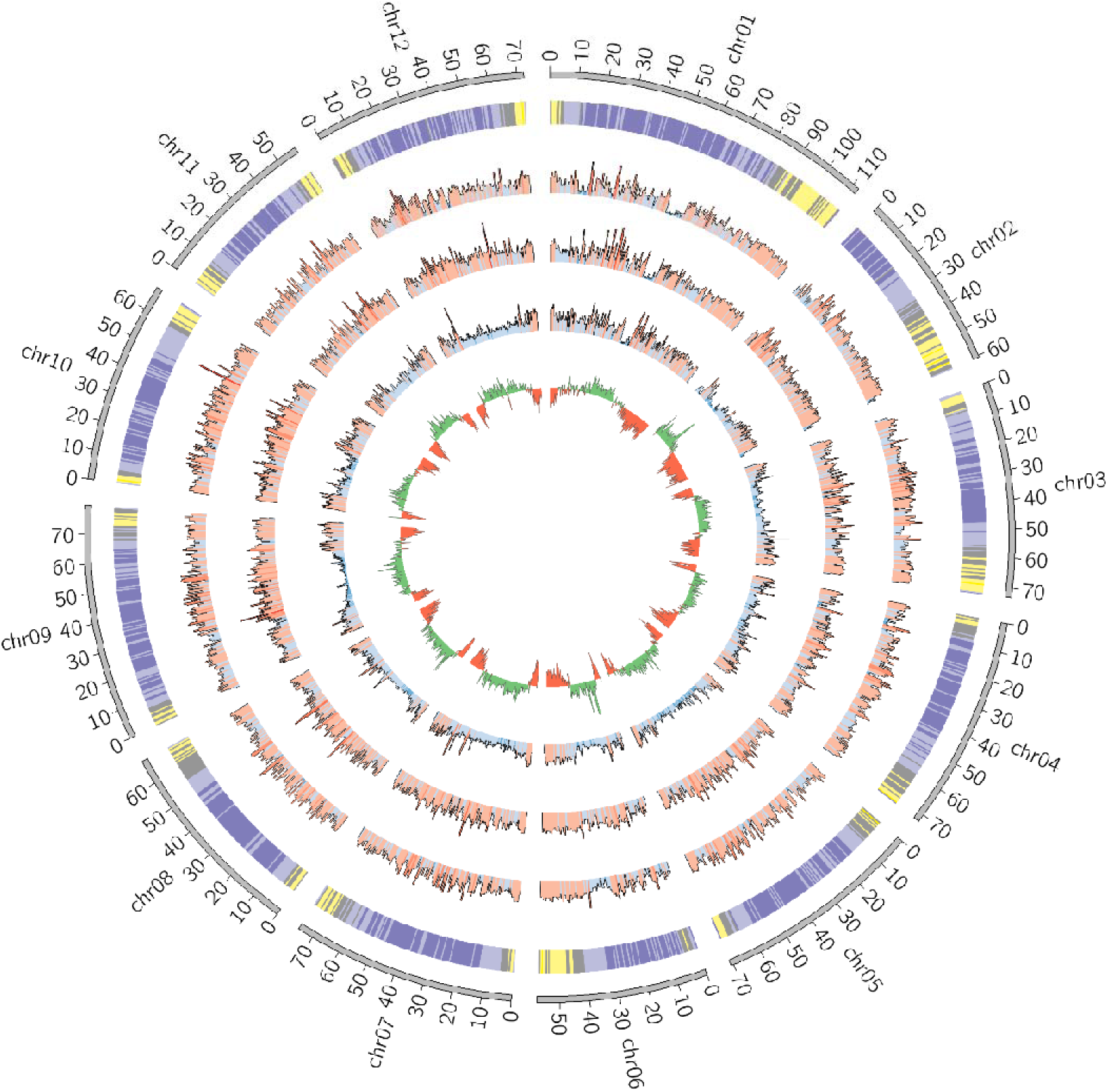
Circos plot of the final S. chilense assembly (scaffolds unmapped to pseudomolecules were omitted). From the outside to the inside, the layers are respectively representing: chromosomes, gene density (purple = low density, yellow = high density), SNP density against S. lycopersicum, SNP density against S. pennellii, SNP density against S. chilense LA3111 (blue = low, red = high) and GC content (green = GC content above the genome mean GC content, red = GC content below the genome mean).

**Table 2:** Statistics of the assembly at each step of the pipeline, obtained with Quast v4.5. The final assembly corresponds to the assembly after removing scaffolds corresponding to organelles and those reported as “to exclude” from the SRA report.

| Stage | Length (Mbp) | # contigs / scaffolds | Largest scaffold (bp) | N50 (bp) |
| --- | --- | --- | --- | --- |
| MaSuRCA (contigs) | 1,001 | 9,780 | 2,713,768 | 190,239 |
| Redundans | 900 | 6,346 | 2,971,814 | 247,906 |
| SSPACE | 913 | 3,066 | 3,721,406 | 489,049 |
| Hybrid Scaffold | 914 | 3,057 | 3,721,406 | 495,590 |
| Arcs + LINKS | 914 | 2,379 | 3,811,166 | 702,252 |
| Pilon | 914 | 2,379 | 3,808,466 | 702,157 |
| Bbmap dedupe | 901 | 1,987 | 3,808,466 | 711,918 |
| GapFiller | 901 | 1,987 | 3,809,012 | 711,918 |
| HiC corrected assembly | 901 | 9,461 | 2,833,999 | 212,100 |
| ALL-HiC + Juicer (chromosomes) | 902 | 12 + 1,906 | 112,267,598 | 72,333,043 |
| **Final assembly** | **901** | **12 + 1,899** | **112,267,598** | **72,333,043** |

The ordering and orientation of the scaffolds with the Hi-C data resulted in a chromosome-level assembly. The assembly has a N50 of 72 Mbp and is composed of 1,911 sequences, corresponding to the 12 chromosomes from *S. chilense* and 1,899 unmapped scaffolds. Notably, 96% of the assembly was found within 12 sequence blocks. The total length of the assembly is 901 Mbp, which is close to both the estimation of 845 Mbp done via the k-mer analysis and the 914 Mbp length from a previously published assembly of the same species, but of a different accession: LA3111 [18]. Figure 5 represents the final contact map, with the pseudomolecules and unplaced scaffolds represented as blue boxes.

**Figure 5:**
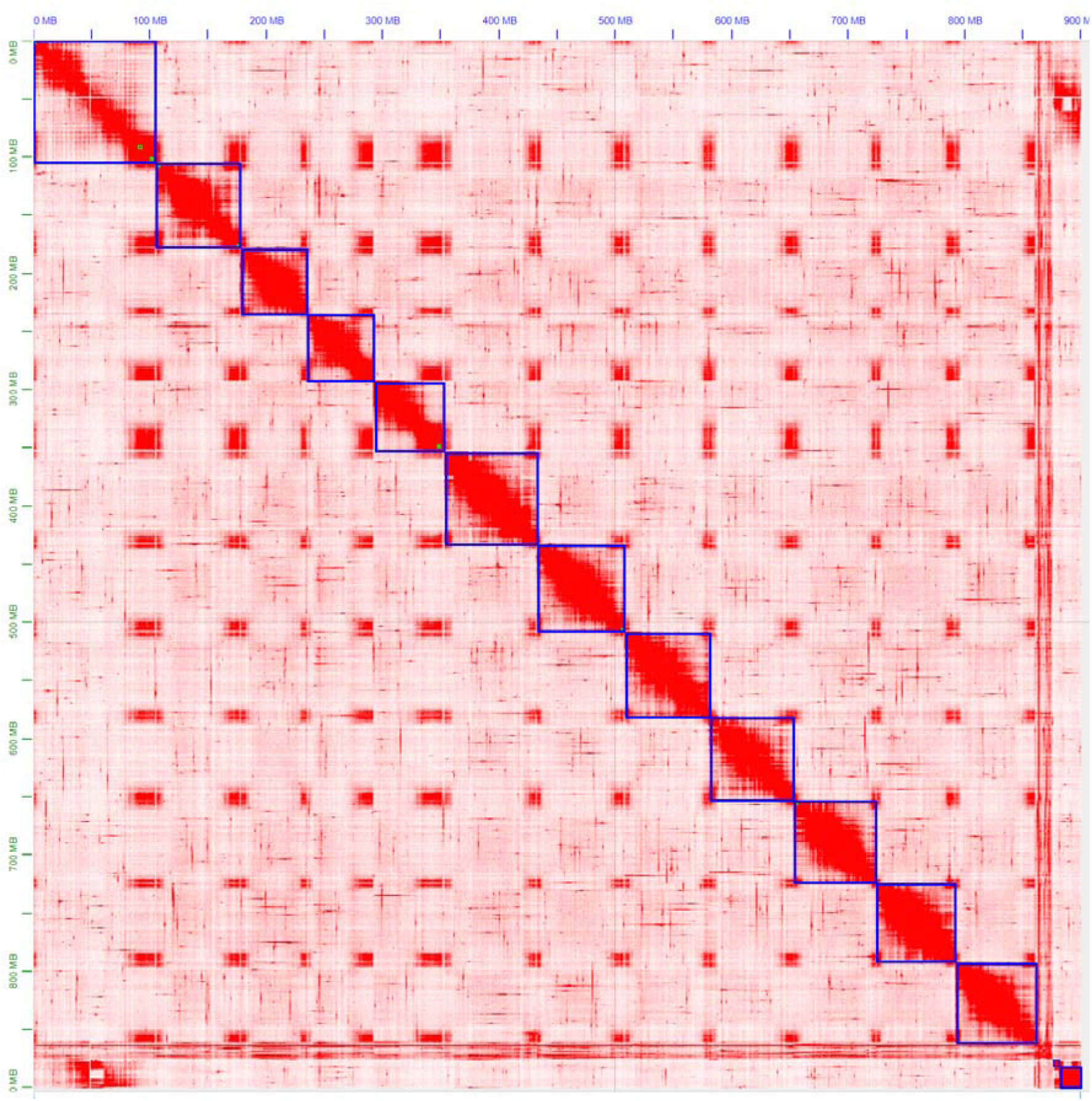
Hi-C map of the final S. chilense (LA1972) assembly, visualised in Juicebox. Hi-C contacts are represented in red, chromosomes and scaffolds are delimited by blue squares. The squares on the bottom right represents unmapped scaffolds.

The assembly was estimated to be 93% complete by KAT, based on a comparison between k-mers obtained from the reads against k-mers from the assembly. This high completeness assessment was confirmed with BUSCO, which managed to identify 95% of the 3,052 orthologues from the *Solanaceae* dataset v10, as complete in the final assembly. These numbers are higher than those obtained from previously published assemblies of wild relatives of tomato, including *S. chilense*, but slightly lower than those obtained from *S. lycopersicum* and *S. pennellii*, which is expected when assembling a heterozygous, self-incompatible wild species. Comparisons of the Quast and BUSCO results with assemblies from other closely related *Solanum* species are available in Table 3 and The BUSCO results obtained at each step of our pipeline are in Supplementary Table 1.

**Table 3:** Comparison of our S. chilense assembly (accession LA1972), against other tomato species. If an assembly is at a chromosome-level, the number of unplaced scaffolds is indicated by the number after the + sign.

| Species | Length (Mbp) | # sequences | Largest scaffold (bp) | N50 (bp) |
| --- | --- | --- | --- | --- |
| *S. chilense* (LA1972) | 901 | 12 + 1,899 | 112,267,598 | 72,333,043 |
| *S. chilense* (LA3111) | 914 | 81,304 | 1,123,112 | 70,632 |
| *S. pennellii* | 990 | 12 + 4,587 | 109,333,515 | 77,991,103 |
| *S. lycopersicum* v4.0 | 783 | 12 + 152 | 90,863,682 | 65,269,487 |
| *S. lycopersicoides* | 1,287 | 12 + 3,084 | 133,548,845 | 93,853,793 |
| *S. pimpinellifolium* | 826 | 107,698 | 893,636 | 78,865 |

**Table 4:** BUSCO results of our S. chilense assembly and other tomato species, based on the Solanaceae dataset v10. (C: Complete, S: Single, D: Duplicated, F: Fragmented, M: Missing BUSCOs)

| Species | Complete [Single, Duplicated], Fragmented and Missing BUSCOs (n = 3,052) |
| --- | --- |
| *S. chilense* (LA1972) | C:94.9% [S:90.8%, D:4.1%], F:1.8%, M:3.3% |
| *S. chilense* (LA3111) | C:90.7% [S:89.5%, D:1.2%], F:4.6%, M:4.7% |
| *S. pennellii* | C:96.8% [S:96.1%, D:0.7%], F:1.3%, M:1.9% |
| *S. lycopersicum* v4.0 | C:95.8% [S:94.9%, D:0.9%], F:1.7%, M:2.5% |
| *S. lycopersicoides* | C:93.3% [S:81.9%, D:11.4%], F:3.6%, M:3.1% |
| *S. pimpinellifolium* | C:93.9% [S:92.1%, D:1.8%], F:2.6%, M:3.5% |

The first iteration of Pilon corrected 164,646 misassemblies, including 109,120 single nucleotide polymorphisms (SNPs) and 55,526 InDels. The corrected assembly was then subjected to a second polishing iteration which corrected 52,653 SNPs and 22,915 InDels. Pilon detected 98.7% of correct bases in the assembly, after the two iterations were performed. The dedupe.sh script from BBMap removed 392 scaffolds, corresponding to 13 Mbp. The largest removed scaffold was 66 kbp long and 98% of the removed scaffolds were smaller than 25 kbp. GapFiller removed 237 gaps, amounting to a total of 23,964 Ns. After the GapFiller step, 12.7 Mbp of unknown bases remained in the assembly, corresponding to ∼1.4% of the assembly length.

Here, we generated the first high-quality, chromosome-level assembly of *S. chilense,* which has comparable or better contiguity and completeness than other wild-relatives of tomato.

### Gene prediction and annotation

RepeatMasker masked 62.63% of the assembly, which is consistent with the repeat content of similar species: 59.5% for *S. pimpinellifolium*, 64% for *S. lycopersicum* [40], 82% for *S. pennellii* [63] and 70% for *S. sitiens* [64].

STAR aligned 95.6% of the RNA-Seq reads to the masked assembly. The hints generated from the alignment allowed Augustus to predict 30,994 gene structures; this number increased to 32,972 when including alternative splicing variants). OmicsBox retrieved blast hits for 30,240 of the genes, including 22,571 annotated with Gene Ontology (GO) terms. The InterProScan search identified 29,934 genes with InterProScan IDs and 18,873 with InterProScan GO terms.

The number of predicted genes is close to those obtained from similar species, 32,273 genes in *S. pennellii,* 34,075 genes in *S. lycopersicum*, 31,164 genes in *S. sitiens*. Stam et al. (2019) found 41,481 genes in *S. chilense* LA3111, including 25,885 high-confidence gene models [65].

### *De novo* transcriptome assembly and annotation

Trinity generated a reference transcriptome assembly consisting of 228,844 transcripts, with a N50 of 2,685 bp and totalling a size of 360 Mbp. The BUSCO analysis identified 83.4% of the expected *Solanaceae* orthologues. After clustering the transcripts with CD-HIT-EST and filtering based on TPM counts lower than 1, the transcriptome was composed of 124,065 transcripts, with a N50 of 2,487 bp and totalling a size of 172 Mbp. The number of complete genes detected by BUSCO remained high, at 83.2%. Moreover, Bowtie2 aligned 98.59% of the RNA-Seq reads back to the assembly, further confirming the completeness of the transcriptome assembly. Of the 124,065 transcripts, 98,361 (79%) had a blast hit and 87,059 (70%) had blast top hits with an e-value ≤ 1×10^-3^ indicating these results to be a strong basis for functional annotation. GO terms were assigned to 64,028 transcripts (52%).

### Genes possessing high impact changes and likely to be involved in abiotic stress tolerance of *Solanum chilense*

*S. chilense* contained 202 genes annotated with the GO terms related to drought, salt, or water (Table 1), including 43 with high impact amino acid variants compared to *S. lycopersicum* proteins (PROVEAN score < -2.5) suggesting functional changes for the selected proteins [60]. These protein variants were reverse translated back to SNPs, based on the protein MAFFT alignments and the nucleotide sequences of the genes from Augustus and ITAG4.1. Tersect, a tool to perform set operations on variant data [66], intersected the resulting SNPs with the 84 publicly available re-sequenced genomes of 12 tomato species [67]. In order to analyse fixed changes in each species, only homozygous variants were considered, which eliminated 5 genes from the total number.

**Error! Reference source not found.** shows the distribution of the impactful PROVEAN variants across the 12 species of tomato represented in the 84 genomes datasets. These variants are representing alleles possessing significant amino acid change in *S. lycopersicum* compared to *S. chilense*, which highlight potential functional shift in these genes in the cultivated tomato. The clustering of the species in the heatmap perfectly matches their belonging to the taxonomic groups of genus *Solanum* Section *Lycopersicon* as defined by Peralta et al. [68], namely the groups Lycopersicon, Arcanum, Eriopersicon and Neolycopersicon. As expected, all 57 variants, from the 38 remaining genes, are present in low percentages in the *S. lycopersicum* accessions. The impactful amino acid changes were further confirmed and reduced with the SIFT4G and PPVED algorithms, the detailed list of the resulting 7 relevant genes with their corresponding variants is shown in Table 5, and are discussed below.

**Table 5:** List of genes with GO terms related to salt and drought, and containing impactful variants (PROVEAN < -2.5, SIFT4G < 0.05 and PPVED > 0.5, marked in bold) in S. lycopersicum compared to S. chilense. “AA change” gives the S. chilense LA1972 amino acid, then the protein position, then the S. lycopersicum Heinz 1706 amino acid (e.g. P63A).

| *S. chilense*<br>gene ID | *S. lycopersicum* ensembl ID<br>(Annotation) | AA<br>change | PROVEAN<br>score | SIFT4G<br>score | PPVED |
| --- | --- | --- | --- | --- | --- |
| g6365.t1 | Solyc03g116610.3,<br>Ethylene-responsive transcription<br>factor WIN1 | P63A | **-7.68** | **0.01** | **0.82** |
| g9938.t1 | Solyc06g060250.3,<br>Aldehyde dehydrogenase | R79W<br>H398Q<br>Y417C | **-3.50**<br>**-7.31**<br>**-8.43** | **0.00**<br>0.37<br>**0.00** | 0.04<br>0.29<br>**0.51** |
| g18960.t1 | Solyc07g065620.5,<br>Poly(A)-specific ribonuclease PARN | S570P<br>A523P | **-4.75**<br>**-3.00** | **0.00**<br>0.41 | **0.90**<br>0.02 |
| g29005.t1 | Solyc08g061140.3,<br>OCP3 | D75G<br>K305R<br>P36L | **-3.98**<br>**-2.69**<br>**-4.87** | 0.12<br>0.17<br>**0.02** | 0.07<br>**0.76**<br>**0.56** |
| g14532.t1 | Solyc09g007870.3,<br>Ethylene-insensitive protein 2 | Q178L<br>S450C<br>H544Q<br>R1194H | **-5.54**<br>**-4.32**<br>**-2.54**<br>**-3.93** | **0.03**<br>**0.01**<br>0.34<br>**0.02** | 0.26<br>0.36<br>0.01<br>**0.85** |
| g14972.t1 | Solyc09g015020.1,<br>class I heat shock protein 3 | S41Y<br>R44P<br>A48D | **-2.67**<br>**-2.79**<br>**-4.78** | **0.02**<br>0.06<br>**0.01** | 0.03<br>0.07<br>**0.68** |
| g27963.t1 | Solyc10g086380.1,<br>GAI-like protein 1 | F271L<br>K295E | **-5.43**<br>**-3.70** | 0.08<br>**0.01** | 0.04<br>**0.79** |

### Solyc03g116610 (Ethylene-responsive transcription factor WIN1)

The *WAX INDUCER 1* (*WIN1*) transcription factor is involved in cuticle biosynthesis in *Arabidopsis thaliana* [69] and its over-expression in tomato from a constitutive promoter improves drought resistance [70] while also decreasing fruit and seed weight [71]. Interestingly, for the variant P63A, the proline is the last amino acid of the AP2 domain conserved across the ERF gene family [71] and is present in the *S. chilense* LA1972 allele and in all accessions of the Arcanum, Eriopersicon and Neolycopersicon groups, but it is replaced by alanine in all the Lycopersicon group accessions and hence is predicted to be deleterious to cuticle development in cultivars (Figure 6). Clearly selection for the *S. chilense* allele might give improved drought resistance via reduced cuticular transpiration [72], but it remains to be seen if this natural variation will also reduce fruit size as occurred with constitutive transgenic over-expression.

**Figure 6:**
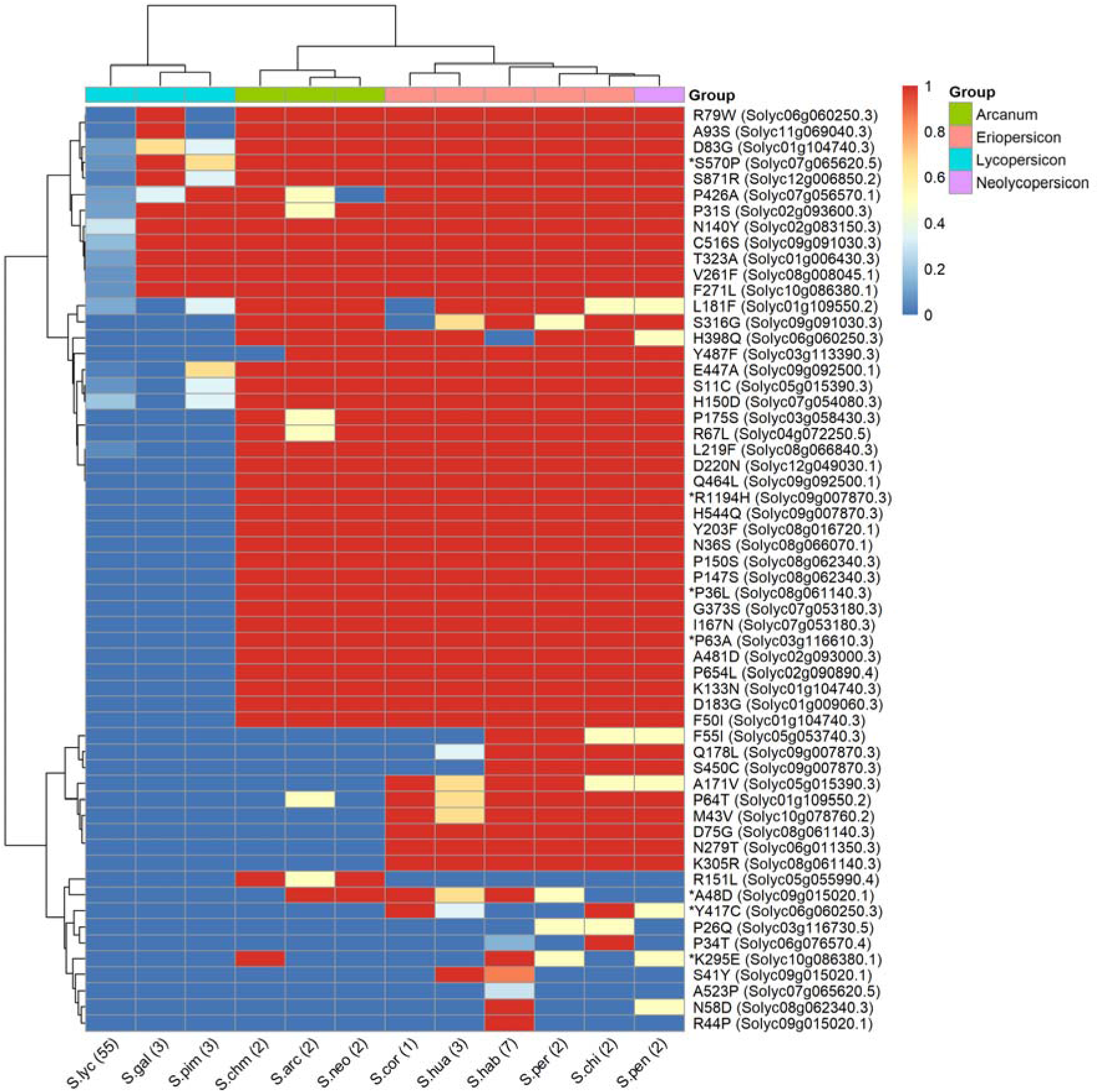
Heatmap showing the similarity of tomato and its related species based on the presence or absence large effect variants within gene with annotation related to salinity and drought stress. Rows are the amino acid changes with a PROVEAN score < -2.5 between S. lycopersicum Heinz 1706 and S. chilense LA1972 as rows, and tomato species from the 84 tomato genomes as columns. The colour represents the proportion (between 0 and 1) of the accessions sharing a common amino acid with S. chilense. The variants, e.g. R79W, are coded as in Table 5. Species were allocated to each group according to taxonomy [96]. The genes identified as impactful by PROVEAN, SIFT4G and PPVED are highlighted by an asterisk.

### Solyc09g007870 (Ethylene-insensitive protein 2)

For R1194H, the *S. chilense* LA1972 allele shares the arginine with all the Arcanum, Eriopersicon and Neolycopersicon accessions, but histidine is present in all Lycopersicon accessions. The R1194H change is in the CEND part of the *EIN2* protein, close to the nuclear localization signal domain [73], which directs this C-terminal part to promote gene expression changes in the nucleus [74]. *EIN2* is a large, complex protein and a component of the intracellular ethylene signalling pathway that stimulates salt tolerance in *A. thaliana* [75], and Solyc09g007870 has been proposed as a candidate gene for a QTL for rootstock-conferred drought resistance [76]. Solyc09g007870 is also involved in regulating tomato fruit ripening and carotenoid accumulation [77]: a large InDel in the promoter of Solyc09g007870 reduced gene expression and was the cause of the yellow-fruited tomato 1 mutation arising in *S. pimpinellifolium* LA1585 [78]. The members of the Lycopersicon group (histidine) have orange or red fruit, while the members of the Arcanum, Eriopersicon and Neolycopersicon groups (arginine) have green fruits [79]. Thus, the histidine variant may have been selected for during evolution of the Lycopersicon group, and domestication of tomato cultivars to provide coloured fruit; it is conceivable this may have been accompanied by a loss of resistance to drought or salinity.

### Solyc06g060250 (aldehyde dehydrogenase; ALDH)

For the Y417C variant, the tyrosine found in *S. chilense* LA1972 is only shared with other accessions of *S. chilense* and accessions from *S. corneliomulleri, S. huaylasense,* and *S. pennellii*; all other accessions have a cysteine. The top blast hit of this gene corresponds to *aldehyde dehydrogenase family 3 member H1* of *A. thaliana*, which is highly expressed upon dehydration, in high salinity stress and under treatment with abscisic acid [80]. The proposed function of stress-responsive ALDH3 family members is the detoxification of aldehydes that accumulate under stress as a result of lipid peroxidation; overexpression of various ALDH genes led to drought and salinity resistance [81]. Y417C is closely located to the protein C-terminus, which is responsible for its dimerization or tetramerization process [82] and thus might alter ALDH enzyme function.

### Solyc10g086380 (a GRAS transcription factor, DELLA subfamily, SlGLD1)

The *S. chilense* LA1972 allele contains two high impact variants which are also present in *S. chilense* LA3111 [65], but absent from the other two *S. chilense* accessions of the 84 tomato genomes dataset (CGN15530 and CGN15532). The Arabidopsis orthologues are involved in the gibberellic acid-mediated signalling and regulation of growth under environmental stresses, including drought [83] and cold [84] and over-expression of *SlGLD1* in tomato gave dwarf plants (J. Li et al., 2015) suggesting the gene is involved in the stress-mediated inhibition of plant growth. The *S. pennellii* LA0716 allele of *SlGLD1* was previously noted to be truncated and inactive due to InDels [85].

### Solyc09g015020 (class I heat shock protein 3 / SlHSP17.7B)

The overexpression of the most homologous Arabidopsis gene, *AtHSP17.8,* in lettuce resulted in dehydration and salt stress resistance phenotypes [86]. However, in tomato *SlHSP17.7B* is expressed specifically during fruit ripening, with low expression in vegetative tissues [87], so there is little evidence for a role in stress tolerance in tomato.

### Solyc07g065620 (poly(A)-specific ribonuclease PARN)

A *PARN* gene in *Arabidopsis thaliana* is responsible for the regulation of appropriate status of poly(A) tract of mitochondrial mRNA [88] and is required for normal ABA, salicylic acid, high salinity and osmotic stress responses [89]. The *S. chilense* LA1972 allele of the S570P variant (serine) is present in all the accessions except some *S. lycopersicum* and *S. pimpinellifolium.* Although the S570P amino acid change appears to be outside the poly(A) polymerase (PAP) domain [90], the rest of the protein is highly conserved, and the amino acid change may still be significant.

### Solyc08g061140 (Homeobox transcription factor / OVEREXPRESSOR OF CATIONIC PEROXIDASE 3; OCP3)

This gene controls an ABA-dependent drought resistance phenotype in Arabidopsis [91], and also mediates resistance to infection by pathogens such as *Botrytis* and *Plectosphaerella* species [92]. The *S. chilense* LA1972 allele is shared with all the accessions of the Arcanum, Eriopersicon and Neolycopersicon groups, but is absent in the Lycopersicon group. The P36L variant is close to RNA polymerase sigma factor domain subunit [90] and might perturb protein function.

### Resources to exploit genetic diversity

Mechanisms for abiotic stress resistance have evolved in wild relatives and their genetics is usually complex and quantitative; these traits can be captured for crop production, for example in land races selected under local sub-optimal environments. However, when modern breeders focus on yield, quality and disease resistance in near optimal conditions, there may be erosion and bottlenecking of genetic variation for abiotic stress resistance [93], and a need to actively introduce natural variation from wild relatives [94, 95] with the support of genetic and genomic resources. To facilitate this, we have created a high-quality, chromosome-scale assembly and annotation of *S. chilense* (LA1972) using a range of sequencing technologies and analysis tools. We are now creating a library of introgressions derived from *S. chilense* LA1972 using the cultivar Kashi Amrit as the genetic background. This combination of genomic and genetic resources will underpin future work to understand and exploit natural genetic variation in this wild relative, and, as a first step, we have used the assembly and annotation to identify amino acid variants present in *S. chilense* LA1972 that could be targets for functional analysis and exploitation in breeding for improved drought and salt resistance. Our analysis highlighted two examples from the literature where there could be counter selection for drought or salinity resistance through pleiotropy: the “dual role” of *WIN* gene (P63A variant) which impacts drought resistance and fruit size; and the R1194H variant of *EIN2*, a gene known to influence both salinity tolerance and fruit colour.

## Supporting information

PacBio Sequel sequence length

Pacbio RSII sequence length

Organille genome assembly

Python script to extract variants from MSA

## Glossary

AA: Amino Acid
ABA: Abscisic Acid
bp: Base pair
BUSCO: Benchmarking Universal Single-Copy Orthologs
GO: Gene Ontology
InDel: Insertion or deletion mutation
IR: Inverted Repeat region
KAT: K-mer Analysis Toolkit
LSC: Large Single Copy region
ORFs: Open Reading Frames
rRNA: Ribosomal RNA
SNP: Single Nucleotide Polymorphism
SSC: Small Single Copy region
TPM: Transcripts Per Million
tRNA: Transfer RNA

## Data availability

*S. chilense* (LA1972) raw sequencing, transcriptome and genome assembly have been deposited at the NCBI’s Sequence Read Archive, under the BioProject number ‘PRJNA880259’.

This Whole Genome Shotgun project has been deposited at DDBJ/ENA/GenBank under the accession JAPDHL000000000. The version described in this paper is version JAPDHL010000000.

The annotated list of predicted genes in GFF3 format, the organelle assemblies and all the commands and scripts used to generate the assembly are available at the following GitHub repository: https://github.com/MCorentin/Solanum_chilense_assembly

## Acknowledgements

We would like to thank Björn Usadel (RWTH Aachen University, Germany) and Richard Finkers and Anthony Bolger (Wageningen University and Research, The Netherlands) for the useful advice and discussions throughout the assembly development. We thank the Earlham Institute and Arima Genomics for providing DNA and RNA sequencing services and the TGRC for providing seeds for *S. chilense* LA1972.

## Author contributions

CM performed most of the genome assembly work, together with TJK (short-reads), DJS,SPK (functional annotation), and JUI (mitochondrial genome) and wrote the initial paper draft, PFA collected the data, AJT designed the experimental analysis. FRM designed the sequencing strategy and supervised the overall assembly approach. All authors contributed equally to reviewing the manuscript.

## References

1. Newman, G., Chapter 22 - Fruit and vegetables: prevention and cure?, in A Prescription for Healthy Living, E. Short, Editor. 2021, Academic Press. p. 243–253.

2. Tomato Genome, C., The tomato genome sequence provides insights into fleshy fruit evolution. Nature, 2012. 485(7400): p. 635–41.

3. Tomato Genome Sequencing, C., et al., Exploring genetic variation in the tomato (Solanum section Lycopersicon) clade by whole-genome sequencing. Plant J, 2014. 80(1): p. 136–48.

4. Zhou, F. and E. Pichersky, The complete functional characterisation of the terpene synthase family in tomato. New Phytol, 2020. 226(5): p. 1341–1360.

5. Gao, L., et al., The tomato pan-genome uncovers new genes and a rare allele regulating fruit flavor. Nat Genet, 2019. 51(6): p. 1044–1051.

6. Alonge, M., et al., Major Impacts of Widespread Structural Variation on Gene Expression and Crop Improvement in Tomato. Cell, 2020. 182(1): p. 145–161 e23.

7. Zhou, Y., et al., Graph pangenome captures missing heritability and empowers tomato breeding. Nature, 2022. 606(7914): p. 527–534.

8. Wang, X., et al., Genome of Solanum pimpinellifolium provides insights into structural variants during tomato breeding. Nat Commun, 2020. 11(1): p. 5817.

9. Molitor, C., et al., De Novo Genome Assembly Of Solanum Sitiens Reveals Structural Variation Associated With Drought And Salinity Tolerance. Bioinformatics, 2021.

10. Stadler, T., U. Arunyawat, and W. Stephan, Population genetics of speciation in two closely related wild tomatoes (Solanum section Lycopersicon). Genetics, 2008. 178(1): p. 339–50.

11. Moyle, L.C., Ecological and evolutionary genomics in the wild tomatoes (Solanum sect. Lycopersicon). Evolution, 2008. 62(12): p. 2995–3013.

12. Chetelat, et al., Distribution, ecology and reproductive biology of wild tomatoes and related nightshades from the Atacama Desert region of northern Chile. Euphytica, 2008. 167(1): p. 77–93.

13. Nakazato, T., D.L. Warren, and L.C. Moyle, Ecological and geographic modes of species divergence in wild tomatoes. American Journal of Botany, 2010. 97(4): p. 680–693.

14. Bigot, S., et al., Comparison of the salt resistance of Solanum lycopersicum x Solanum chilense hybrids and their parents. Frontiers in Horticulture, 2023. 2.

15. Chetelat, R.T., et al., Distribution, ecology and reproductive biology of wild tomatoes and related nightshades from the Atacama Desert region of northern Chile. Euphytica, 2009. 167(1): p. 77–93.

16. Ji, Y., D.J. Schuster, and J.W. Scott, Ty-3, a begomovirus resistance locus near the Tomato yellow leaf curl virus resistance locus Ty-1 on chromosome 6 of tomato. Molecular Breeding, 2007. 20(3): p. 271–284.

17. Böndel, K.B., et al., North–South Colonization Associated with Local Adaptation of the Wild Tomato Species Solanum chilense. Molecular Biology and Evolution, 2015. 32(11): p. 2932–2943.

18. Stam, R., et al., The de Novo Reference Genome and Transcriptome Assemblies of the Wild Tomato Species Solanum chilense Highlights Birth and Death of NLR Genes Between Tomato Species. G3: Genes, Genomes, Genetics, 2019. 9(12): p. 3933–3941.

19. Rick, C.M., Potential genetic resources in tomato species: clues from observations in native habitats. Basic Life Sci, 1973. 2: p. 255–69.

20. Stam, R., et al., The de Novo Reference Genome and Transcriptome Assemblies of the Wild Tomato Species Solanum chilense Highlights Birth and Death of NLR Genes Between Tomato Species. G3 (Bethesda), 2019. 9(12): p. 3933–3941.

21. Yeo, S., et al., ARCS: scaffolding genome drafts with linked reads. Bioinformatics, 2018. 34(5): p. 725–731.

22. Song, L. and L. Florea, Rcorrector: efficient and accurate error correction for Illumina RNA-seq reads. GigaScience, 2015. 4(1): p. 48.

23. Bioinformatics, B. Trim Galore!

24. Martin, M., Cutadapt removes adapter sequences from high-throughput sequencing reads. EMBnet.journal, 2011. 17(1): p. 10–12.

25. Williams, D., et al., Rapid quantification of sequence repeats to resolve the size, structure and contents of bacterial genomes. BMC Genomics, 2013. 14: p. 537.

26. Marçais, G. and C. Kingsford, A fast, lock-free approach for efficient parallel counting of occurrences of k-mers. Bioinformatics, 2011. 27(6): p. 764–770.

27. Zimin, A.V., et al., The MaSuRCA genome assembler. Bioinformatics, 2013. 29(21): p. 2669–2677.

28. Manni, M., et al., BUSCO Update: Novel and Streamlined Workflows along with Broader and Deeper Phylogenetic Coverage for Scoring of Eukaryotic, Prokaryotic, and Viral Genomes. Molecular Biology and Evolution, 2021. 38(10): p. 4647–4654.

29. Pryszcz, L.P. and T. Gabaldón, Redundans: an assembly pipeline for highly heterozygous genomes. Nucleic Acids Research, 2016. 44(12): p. e113–e113.

30. Boetzer, M., et al., Scaffolding pre-assembled contigs using SSPACE. Bioinformatics, 2011. 27(4): p. 578–579.

31. Shelton, J.M., et al., Tools and pipelines for BioNano data: molecule assembly pipeline and FASTA super scaffolding tool. BMC Genomics, 2015. 16(1): p. 734.

32. Warren, R.L., et al., LINKS: Scalable, alignment-free scaffolding of draft genomes with long reads. GigaScience, 2015. 4(1): p. 35.

33. Li, H. and R. Durbin, Fast and accurate short read alignment with Burrows–Wheeler transform. Bioinformatics, 2009. 25(14): p. 1754–1760.

34. Walker, B.J., et al., Pilon: An Integrated Tool for Comprehensive Microbial Variant Detection and Genome Assembly Improvement. PLOS ONE, 2014. 9(11): p. e112963.

35. Li, H., et al., The Sequence Alignment/Map format and SAMtools. Bioinformatics, 2009. 25(16): p. 2078–2079.

36. Nadalin, F., F. Vezzi, and A. Policriti, GapFiller: a de novo assembly approach to fill the gap within paired reads. BMC Bioinformatics, 2012. 13(14): p. S8.

37. Bolger, A.M., M. Lohse, and B. Usadel, Trimmomatic: a flexible trimmer for Illumina sequence data. Bioinformatics, 2014. 30(15): p. 2114–2120.

38. Burton, J.N., et al., Chromosome-scale scaffolding of de novo genome assemblies based on chromatin interactions. Nature Biotechnology, 2013. 31(12): p. 1119–1125.

39. Zhang, X., et al., Assembly of allele-aware, chromosomal-scale autopolyploid genomes based on Hi-C data. Nature Plants, 2019. 5(8): p. 833–845.

40. Hosmani, P.S., et al., An improved de novo assembly and annotation of the tomato reference genome using single-molecule sequencing, Hi-C proximity ligation and optical maps. 2019.

41. Bolger, A., et al., The genome of the stress-tolerant wild tomato species Solanum pennellii. Nature Genetics, 2014. 46(9): p. 1034–1038.

42. Durand, N.C., et al., Juicebox Provides a Visualization System for Hi-C Contact Maps with Unlimited Zoom. Cell Systems, 2016. 3(1): p. 99–101.

43. Gurevich, A., et al., QUAST: quality assessment tool for genome assemblies. Bioinformatics, 2013. 29(8): p. 1072–1075.

44. Kriventseva, E.V., et al., OrthoDB v10: sampling the diversity of animal, plant, fungal, protist, bacterial and viral genomes for evolutionary and functional annotations of orthologs. Nucleic Acids Research, 2019. 47(D1): p. D807–D811.

45. Marçais, G., et al., MUMmer4: A fast and versatile genome alignment system. PLOS Computational Biology, 2018. 14(1): p. e1005944.

46. Mapleson, D., et al., KAT: a K-mer analysis toolkit to quality control NGS datasets and genome assemblies. Bioinformatics, 2017. 33(4): p. 574–576.

47. Stanke, M. and B. Morgenstern, AUGUSTUS: a web server for gene prediction in eukaryotes that allows user-defined constraints. Nucleic acids research, 2005. 33(Web Server issue): p. W465–7.

48. Smit, A., Hubley, R & Green, P. RepeatMasker Open-4.0. 2013 2013–2015.

49. Fernandez-Pozo, N., et al., The Sol Genomics Network (SGN)—from genotype to phenotype to breeding. Nucleic Acids Research, 2015. 43(Database issue): p. D1036–D1041.

50. Dobin, A., et al., STAR: ultrafast universal RNA-seq aligner. Bioinformatics, 2013. 29(1): p. 15–21.

51. Conesa, A. and S. Götz, Blast2GO: A comprehensive suite for functional analysis in plant genomics. International journal of plant genomics, 2008. 2008: p. 619832.

52. Jones, P., et al., InterProScan 5: genome-scale protein function classification. Bioinformatics, 2014. 30(9): p. 1236–1240.

53. Grabherr, M.G., et al., Trinity: reconstructing a full-length transcriptome without a genome from RNA-Seq data. Nature biotechnology, 2011. 29(7): p. 644–652.

54. Li, W. and A. Godzik, Cd-hit: a fast program for clustering and comparing large sets of protein or nucleotide sequences. Bioinformatics, 2006. 22(13): p. 1658–1659.

55. Langmead, B. and S.L. Salzberg, Fast gapped-read alignment with Bowtie 2. Nature Methods, 2012. 9(4): p. 357–359.

56. Altschul, S.F., et al., Basic local alignment search tool. Journal of Molecular Biology, 1990. 215(3): p. 403–410.

57. Berardini, T.Z., et al., The Arabidopsis Information Resource: Making and Mining the ‘Gold Standard’ Annotated Reference Plant Genome. Genesis (New York, N.Y. : 2000), 2015. 53(8): p. 474–485.

58. Shen, W., et al., SeqKit: A Cross-Platform and Ultrafast Toolkit for FASTA/Q File Manipulation. PLoS ONE, 2016. 11(10): p. e0163962.

59. Katoh, K., J. Rozewicki, and K.D. Yamada, MAFFT online service: multiple sequence alignment, interactive sequence choice and visualization. Briefings in Bioinformatics, 2019. 20(4): p. 1160–1166.

60. Choi, Y., et al., Predicting the Functional Effect of Amino Acid Substitutions and Indels. PLoS ONE, 2012. 7(10): p. e46688.

61. Vaser, R., et al., SIFT missense predictions for genomes. Nature Protocols, 2016. 11(1): p. 1–9.

62. Gou, X., et al., PPVED: A machine learning tool for predicting the effect of single amino acid substitution on protein function in plants. Plant Biotechnology Journal, 2022. 20(7): p. 1417–1431.

63. Bolger, A., et al., The genome of the stress-tolerant wild tomato species Solanum pennellii. Nature Genetics, 2014. 46(9): p. 1034–1038.

64. Molitor, C., et al., De Novo Genome Assembly Of Solanum Sitiens Reveals Structural Variation Associated With Drought And Salinity Tolerance. Bioinformatics, 2021(btab048).

65. Stam, R., et al., The de Novo Reference Genome and Transcriptome Assemblies of the Wild Tomato Species Solanum chilense Highlights Birth and Death of NLR Genes Between Tomato Species. G3 (Bethesda, Md.), 2019. 9(12): p. 3933–3941.

66. Kurowski, T.J. and F. Mohareb, Tersect: a set theoretical utility for exploring sequence variant data. Bioinformatics, 2020. 36(3): p. 934–935.

67. Consortium, T.G.S., et al., Exploring genetic variation in the tomato (Solanum section Lycopersicon) clade by whole-genome sequencing. The Plant Journal: For Cell and Molecular Biology, 2014. 80(1): p. 136–148.

68. Peralta, I.E., D.M. Spooner, and S. Knapp, Taxonomy of Wild Tomatoes and Their Relatives (Solanum sect. Lycopersicoides, sect. Juglandifolia, sect. Lycopersicon; Solanaceae). Systematic Botany Monographs, 2008. 84: p. 1–186.

69. Aharoni, A., et al., The SHINE clade of AP2 domain transcription factors activates wax biosynthesis, alters cuticle properties, and confers drought tolerance when overexpressed in Arabidopsis. The Plant Cell, 2004. 16(9): p. 2463–2480.

70. Al-Abdallat, A.M., et al., Over-Expression of SlSHN1 Gene Improves Drought Tolerance by Increasing Cuticular Wax Accumulation in Tomato. International Journal of Molecular Sciences, 2014. 15(11): p. 19499–19515.

71. Li, Q., et al., Differential expression of SlKLUH controlling fruit and seed weight is associated with changes in lipid metabolism and photosynthesis-related genes. Journal of Experimental Botany, 2021. 72(4): p. 1225–1244.

72. Boyer, J.S., Turgor and the transport of CO2 and water across the cuticle (epidermis) of leaves. Journal of Experimental Botany, 2015. 66(9): p. 2625–2633.

73. Wen, X., et al., Activation of ethylene signaling is mediated by nuclear translocation of the cleaved EIN2 carboxyl terminus. Cell Research, 2012. 22(11): p. 1613–1616.

74. Zhang, J., et al., Uncertainty of EIN2Ser645/Ser924 Inactivation by CTR1-Mediated Phosphorylation Reveals the Complexity of Ethylene Signaling. Plant Communications, 2020. 1(3): p. 100046.

75. Lei, G., et al., EIN2 regulates salt stress response and interacts with a MA3 domain-containing protein ECIP1 in Arabidopsis. Plant, Cell & Environment, 2011. 34(10): p. 1678–1692.

76. Asins, M.J., et al., Genetic Analysis of Root-to-Shoot Signaling and Rootstock-Mediated Tolerance to Water Deficit in Tomato. Genes, 2021. 12(1): p. 10.

77. Karlova, R., et al., Transcriptional control of fleshy fruit development and ripening. Journal of Experimental Botany, 2014. 65(16): p. 4527–4541.

78. Gao, L., et al., The yellow-fruited tomato 1 (yft1) mutant has altered fruit carotenoid accumulation and reduced ethylene production as a result of a genetic lesion in ETHYLENE INSENSITIVE2. Theoretical and Applied Genetics, 2016. 129(4): p. 717–728.

79. Gonzali, S. and P. Perata, Fruit Colour and Novel Mechanisms of Genetic Regulation of Pigment Production in Tomato Fruits. Horticulturae, 2021. 7(8): p. 259.

80. Kirch, H.-H., et al., Detailed expression analysis of selected genes of the aldehyde dehydrogenase (ALDH) gene superfamily in Arabidopsis thaliana. Plant Molecular Biology, 2005. 57(3): p. 315–332.

81. Stiti, N., V. Giarola, and D. Bartels, From algae to vascular plants: The multistep evolutionary trajectory of the ALDH superfamily towards functional promiscuity and the emergence of structural characteristics. Environmental and Experimental Botany, 2021. 185: p. 104376.

82. Shortall, K., et al., Insights into Aldehyde Dehydrogenase Enzymes: A Structural Perspective. Frontiers in Molecular Biosciences, 2021. 8.

83. Wang, Z., et al., GAI Functions in the Plant Response to Dehydration Stress in Arabidopsis thaliana. International Journal of Molecular Sciences, 2020. 21(3): p. 819.

84. Lantzouni, O., et al., GROWTH-REGULATING FACTORS Interact with DELLAs and Regulate Growth in Cold Stress. The Plant Cell, 2020. 32(4): p. 1018–1034.

85. Alseekh, S., et al., Identification and Mode of Inheritance of Quantitative Trait Loci for Secondary Metabolite Abundance in Tomato. The Plant Cell, 2015. 27(3): p. 485–512.

86. Kim, D.H., Z.-Y. Xu, and I. Hwang, AtHSP17.8 overexpression in transgenic lettuce gives rise to dehydration and salt stress resistance phenotypes through modulation of ABA-mediated signaling. Plant Cell Reports, 2013. 32(12): p. 1953–1963.

87. Upadhyay, R.K., M.L. Tucker, and A.K. Mattoo, Ethylene and RIPENING INHIBITOR Modulate Expression of SlHSP17.7A, B Class I Small Heat Shock Protein Genes During Tomato Fruit Ripening. Frontiers in Plant Science, 2020. 11.

88. Hirayama, T., et al., A poly(A)-specific ribonuclease directly regulates the poly(A) status of mitochondrial mRNA in Arabidopsis. Nature Communications, 2013. 4(1): p. 2247.

89. Nishimura, N., et al., Analysis of ABA Hypersensitive Germination2 revealed the pivotal functions of PARN in stress response in Arabidopsis. The Plant Journal, 2005. 44(6): p. 972–984.

90. Marchler-Bauer, A., et al., CDD/SPARCLE: functional classification of proteins via subfamily domain architectures. Nucleic Acids Research, 2017. 45(D1): p. D200–D203.

91. Ramírez, V., et al., Drought tolerance in Arabidopsis is controlled by the OCP3 disease resistance regulator. The Plant Journal, 2009. 58(4): p. 578–591.

92. Coego, A., et al., An Arabidopsis homeodomain transcription factor, OVEREXPRESSOR OF CATIONIC PEROXIDASE 3, mediates resistance to infection by necrotrophic pathogens. The Plant cell, 2005. 17(7): p. 2123–2137.

93. van de Wouw, M., et al., Genetic erosion in crops: concept, research results and challenges. Plant Genetic Resources-Characterization and Utilization, 2010. 8(1): p. 1–15.

94. Pereira, L., et al., Natural Genetic Diversity in Tomato Flavor Genes. Frontiers in Plant Science, 2021. 12.

95. Kulus, D., Genetic resources and selected conservation methods of tomato. Journal of Applied Botany and Food Quality, 2018. 91: p. 135–144.

96. Peralta, I.E., D.M. Spooner, and S. Knapp, Taxonomy of wild tomatoes and their relatives (Solanum sect. Lycopersicoides, sect. Juglandifolia, sect. Lycopersicon; Solanaceae), in Systematic Botany Monographs. 2008, American Society of Plant Taxonomists. p. 1–186.

