## Supplementary figures and images for "A chromosome-level genome assembly of *Solanum chilense*, a tomato wild relative associated with resistance to salinity and drought"

### Pacbio RSII sequence length

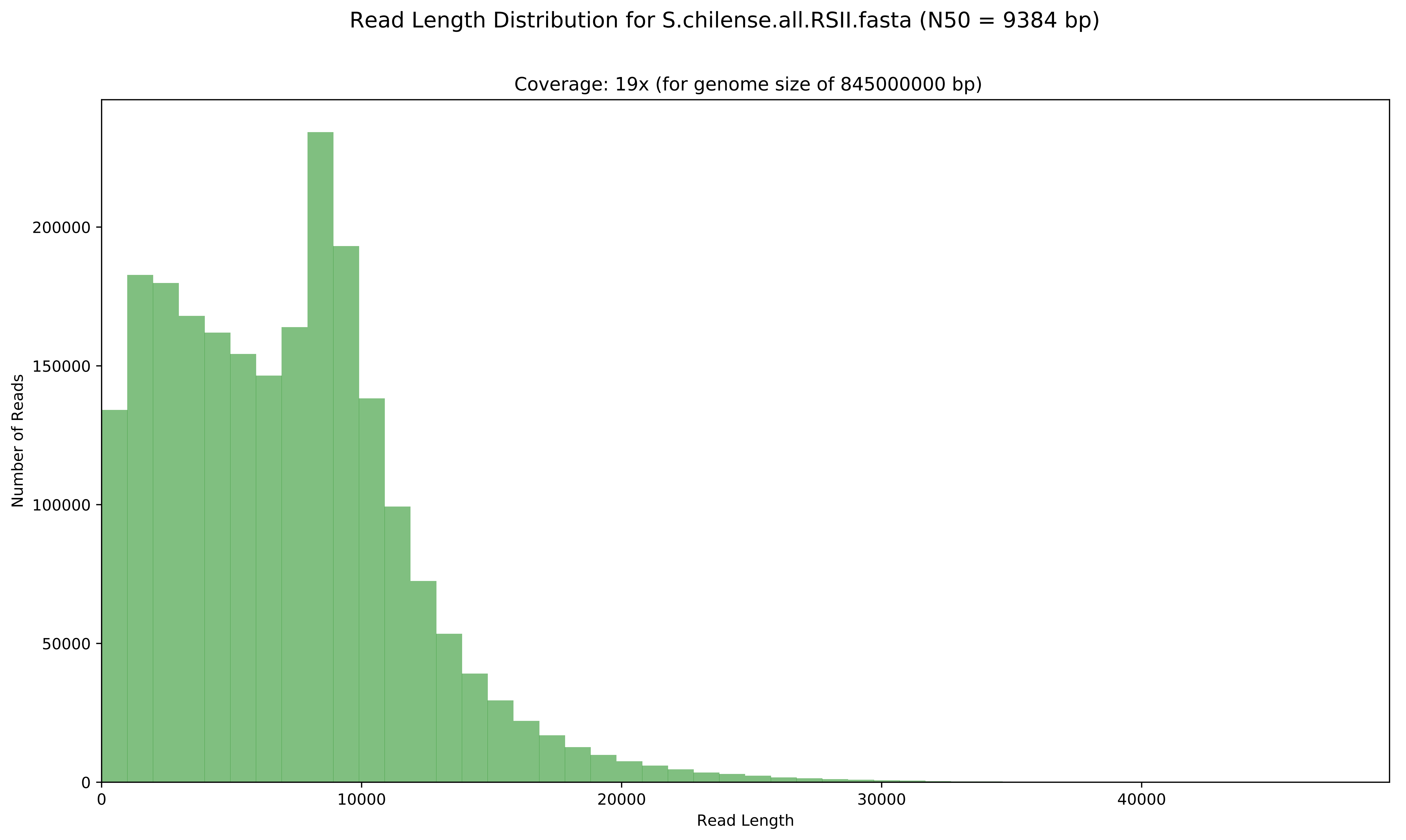

### PacBio Sequel sequence length

#### Read Length Distribution for S.chilense_sequel_ALL.subreads.fasta (N50 = 14770 bp)

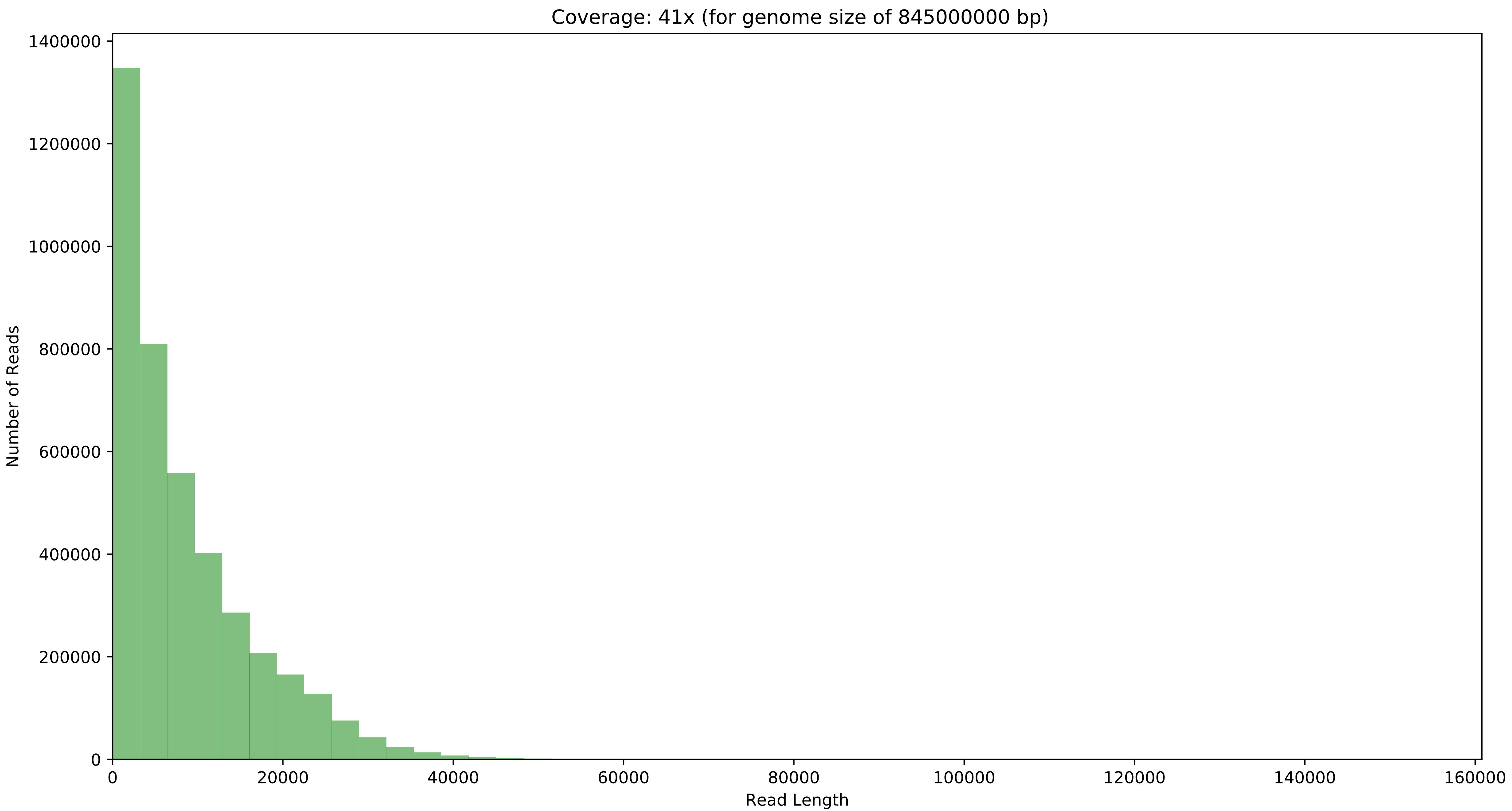
