## Supplementary material for "A chromosome-level genome assembly of *Solanum chilense*, a tomato wild relative associated with resistance to salinity and drought": Organille genome assembly

### Organelles assemblies and annotations

#### Methods

Assembly of the chloroplast genome was performed using getOrganelle v1.7.5 (Jin et al., 2020) using the *get_organelle_from_reads.py* script with the Illumina reads as input and the following parameters: the organelle type (-F) was set to *embplant_pt*, and the maximum number of rounds (-R) was set to 15. First, getOrganelle identified reads pertaining to the chloroplast genome from the Whole Genome library (using a seed database) before proceeding to the assembly with SPAdes (Bankevich et al., 2012).

The mitochondrial genome was assembled with NOVOPlasty v4.3.1 (Dierckxsens et al., 2017) using a k-mer size of 30. The nucleotide sequence of the *NAD3* gene from *S. lycopersicum* was used as a seed to identify the Illumina reads pertaining to the mitochondrial genome, the mitochondrial genome of *S. lycopersicum* (NC_035963.1) was used as the mitochondrial reference and the chloroplast assembly previously produced by getOrganelle was used as the chloroplast reference.

Both the chloroplast and mitochondrial assemblies were given as input to GeSeq web service for annotation (Tillich et al., 2017). GeSeq relies on BLATN, ARAGORN v1.2.38, ARWEN v1.2.3, Chloë v0.1.0 and tRNAscan-SE v2.0.7 to identify chloroplast and/or mitochondrial genes. The resulting annotations were visualised with OGDRAW (Greiner et al., 2019) as Circos plots, (Figure 6, Figure 7).

The chloroplast and mitochondrial assemblies were blasted against the main assembly to identify potential unmapped scaffolds corresponding to organelle sequences. The alignment was performed by *blastn* v2.6.0 (Camacho et al., 2009) with the following parameters: *-perc_identity 95 -max_hsps 2 -outfmt 6 -max_target_seqs 2*. Then, we manually identified relevant blast hits depending on the alignment’s length and percentage of identity. The blast search identified 5 scaffolds with >99% identity with the organelle sequences and containing exclusively organellar genes. The genes corresponding to the removed sequences were detected and then removed from the annotation with *bedtools intersect* and *bedtools subtract*.

#### Results

#### Chloroplast genome

The chloroplast genome was assembled using getOrganelle as a continuous 155,559 bp long sequence, with a GC content of 37.85%. The assembly graph, in *Graphical Fragment Assembly* format, was visualised using Bandage (Wick et al., 2015). The assembly conformed to the typical circular quadripartite structure of chloroplast genomes, composed of a Large Single Copy (LSC) region and a Small Single Copy (SSC) region separated by two copies of the Inverted Repeat (IR) region (Figure 5). The length of each component was similar to that reported for other *Solanum* species: 86 kb for the LSC, 25 kb for each IR and 18 kb for the SSC (D. Li et al., n.d.).

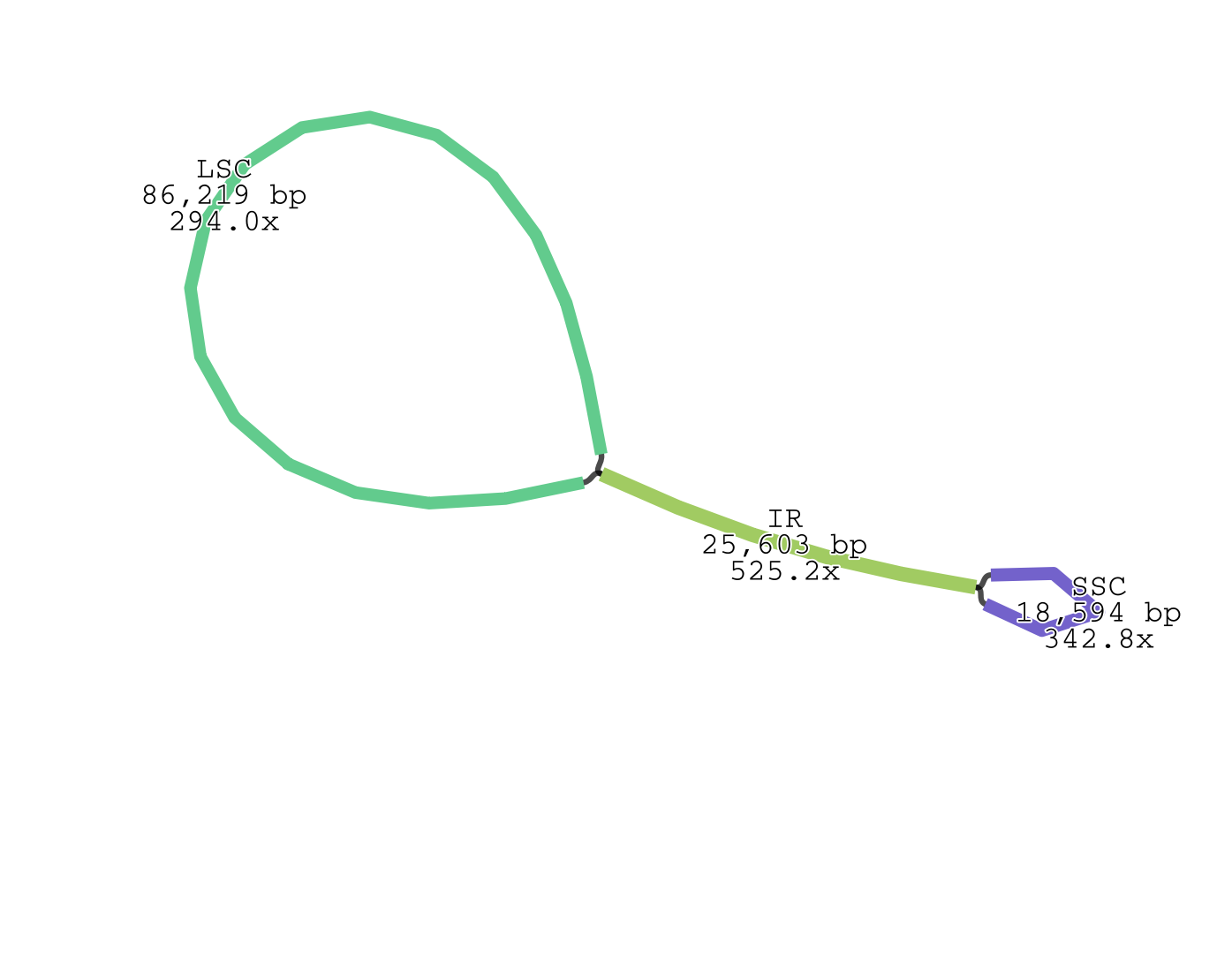

Figure 5: Representation of the graph file generated by getOrganelle for the chloroplast genome of S. chilense, visualised using Bandage. The circular quadripartite conformation is visible, with the Large Single Copy region (SLC), the Small Single Copy region (SSC) and the pair of Inverted Repeat regions (IR).

The annotation from GeSeq identified 116 unique genes in the *S. chilense* chloroplast genome, including 4 ribosomal RNA (rRNA) genes and 31 transfer RNA (tRNA) genes. These numbers are also similar to other *Solanum* species (D. Li et al., n.d.). The chloroplast assembly and its annotation are represented as a Circos plot in Figure 6.

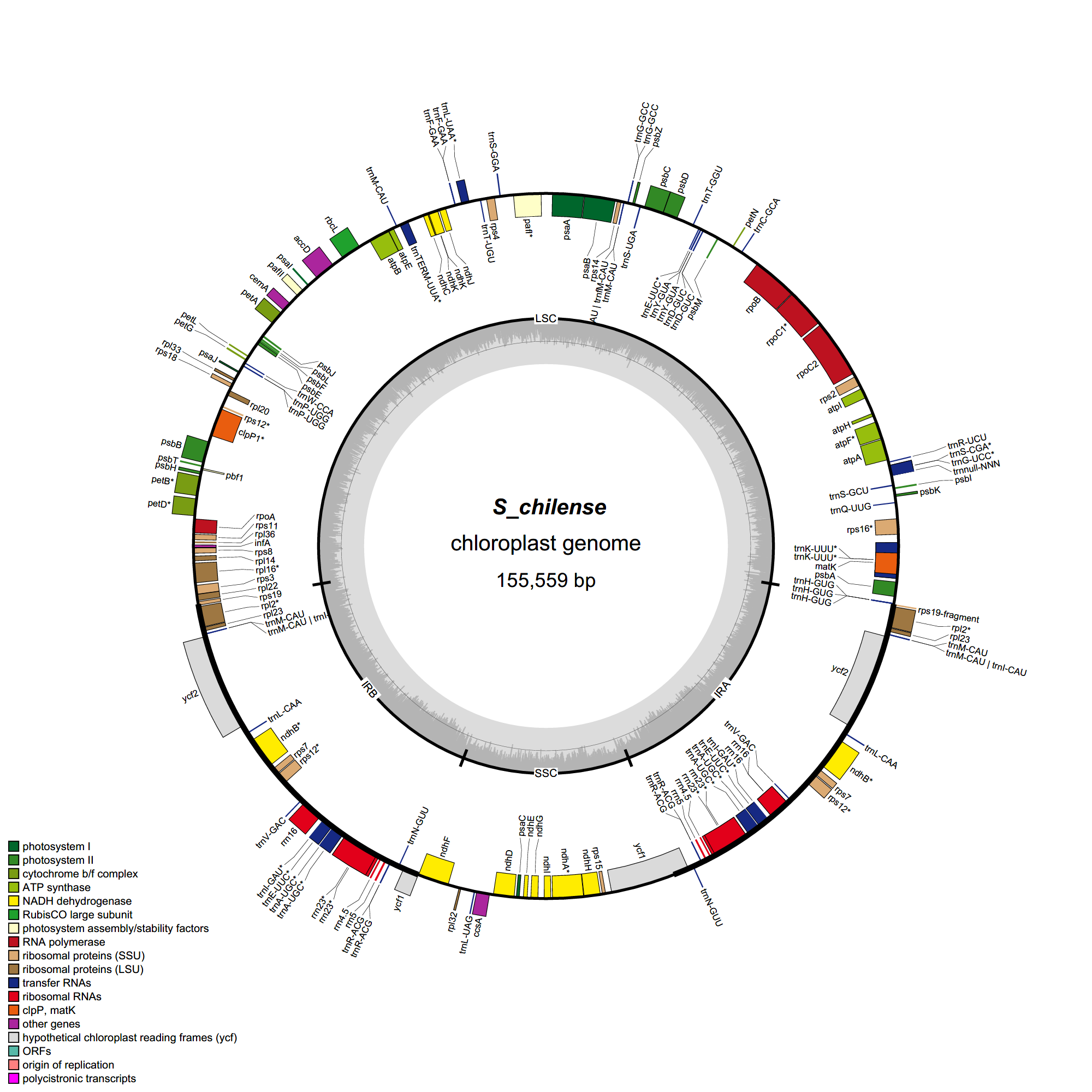

Figure 6: Chloroplast genome of S. chilense. The inner circle represents the GC content, with the middle line marking the 50% threshold. Genes inside the circle are transcribed clockwise, genes outside the circle counterclockwise. Intron containing genes are marked by an asterisk (*). LSC, Large Single Copy region; SSC, Small Single Copy region; IRA and IRB, Inverted Repeat A and B, respectively.

#### Mitochondrial genome

NOVOPlasty assembled the mitochondrial genome into a single circularised sequence of 446,493 bp. This is within the expected range of *Solanum* mitochondrial genomes: a recent study by Kim and Lee (H. T. Kim & Lee, 2018) assembled three mitochondrial genomes, two of *S. lycopersicum* and one of *S. pennellii*, with lengths varying from 423,596 to 446,257 bp. The annotation from GeSeq identified 82 unique genes, including 3 rRNAs and 50 tRNAs. A representation of the mitochondrial assembly with its annotation as a Circos plot is available in Figure 7.

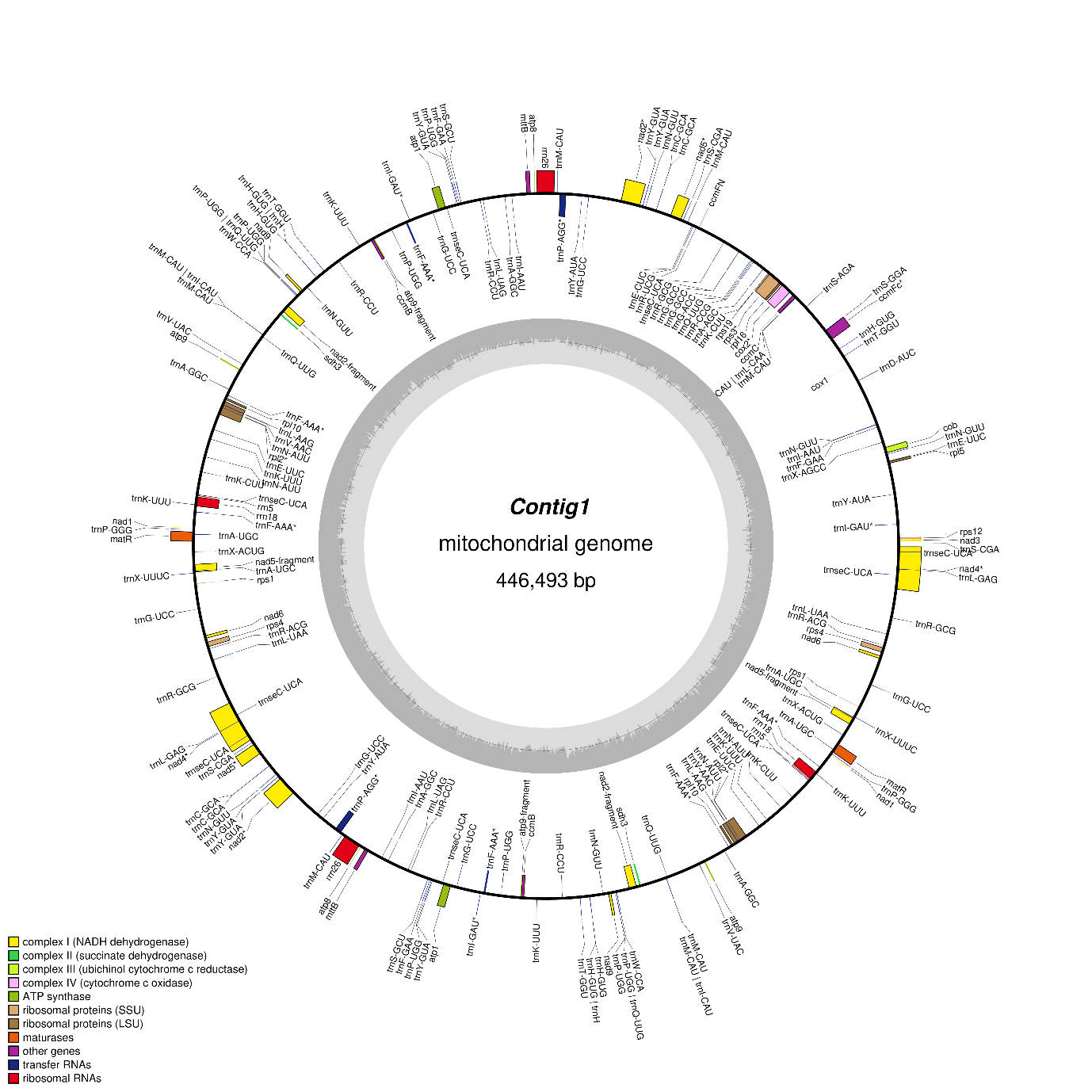

Figure 7: Mitochondrial genome of S. chilense. The inner circle represents the GC content, with the middle line marking the 50% threshold. Genes inside the circle are transcribed clockwise, genes outside the circle counter clockwise. Intron containing genes are marked by an asterisk (*).
