## Supplementary material for "A chromosome-level genome assembly of *Solanum chilense*, a tomato wild relative associated with resistance to salinity and drought": Python script to extract variants from MSA

```
import sys
import os
import getopt
from Bio import SeqIO


fasta = None


def usage():
    print("\nUsage:")
    print("python extract_variants_indels_from_MSA.py -f
input.fasta\n")
    print("-f input.fasta: the input fasta file, resulting from a
Multiple Sequence Alignment. (For now this script only accepts
alignments between two sequences)")
    print("-h: print the help and exit")


try:
    opts, args = getopt.getopt(sys.argv[1:], "hf:", ["help",
"fasta="])
except getopt.GetoptError as err:
    # Print an error message if the option is not recognized:
    print(str(err))
    usage()
    sys.exit(2)

for o, a in opts:
    if o in ("-f", "--fasta"):
        fasta = a
    elif o in ("-h", "--help"):
        usage()
        sys.exit()
    else:
        assert False, "unhandled option"

if(fasta == None):
    print("Please provide a fasta file as input (option -f)")
    usage()
    sys.exit()

if not(os.path.exists(fasta)):
    print(str(fasta) + " does not exists !\n")
    sys.exit()


seq_records = SeqIO.parse(fasta, 'fasta')
# We only support fasta files with two sequences for now:
refseq_record = next(seq_records)
seq_record = next(seq_records)
# The for loop is based on an iterator because we need to use
"next()" during the
# while loops for the insertions and deletions.
```

```
index_iter = iter(range(0, len(refseq_record)))
# ref_real_pos used to keep track of the actual position of the
nucleotides in the
# gene. Indeed, in the MAFFT file, there are some "-" characters for
the insertions
# and deletions. We need to "skip" them when getting the positions
of the variants:
ref_real_pos = 1

# We check one nucleotide at a time from the reference, since it's a
MAFFT alingment
# the two sequences should have the same lengths (with insertions /
deletions written
# as "----").
for i in index_iter:
    ref_aa = refseq_record[i]
    seq_aa = seq_record[i]

    # Case: substitution, we record it and update the "real"
positions:
    if(ref_aa != seq_aa and ref_aa != "-" and seq_aa != "-"):
        print(ref_aa + str(ref_real_pos) + seq_aa)

    # If the reference has a "-", then it's an insertion in the
query:
    # We use a while loop to get to the end of the insertion
    if(ref_aa == "-"):
        # If the insertion is at the very start of the gene, then
the format
        # only take the first amino acid, eg: "M1insMYUPT" instead
of "M1_A2insMYUPT"
        if(i > 0):
            start = refseq_record[i-1] + str(ref_real_pos-1) + "_"
        else:
            start = ""

        # We save the inserted nucleotides (in query) inside
"nucleotide":
        insertion = seq_aa
        while(i < (len(refseq_record)-1) and refseq_record[i+1] ==
"-"):
            i = next(index_iter)
            insertion = insertion + seq_record[i]

        # If "i" is equal to the reference length, then the
insertion is at the
        # end, so we remove the "_" from 'start' and we don't need
an 'end':
        if(i == len(refseq_record)-1):
            start = start[0:len(start)-1]
            end = ""
        else:
            end = refseq_record[i+1] + str(ref_real_pos)
```

```
        print(start + end + "ins" + insertion)

    # If the query has a "-", then it is a deletion in the query:
    # We use a while loop to get to the end of the insertion
    if(seq_aa == "-"):
        deletion = ref_aa + str(ref_real_pos)
        while(i < (len(seq_record)-1) and seq_record[i+1] == "-"):
            i = next(index_iter)
            ref_real_pos = ref_real_pos + 1
            deletion = deletion + "_" + refseq_record[i] +
str(ref_real_pos)
        print(deletion + "del")

    if(refseq_record[i] != "-"):
        ref_real_pos = ref_real_pos + 1
```
